# Hippocampal sharp wave-ripples and the associated sequence replay emerge from structured synaptic interactions in a network model of area CA3

**DOI:** 10.1101/2021.02.18.431868

**Authors:** András Ecker, Bence Bagi, Eszter Vértes, Orsolya Steinbach-Németh, Mária R. Karlócai, Orsolya I. Papp, István Miklós, Norbert Hájos, Tamás F. Freund, Attila I. Gulyás, Szabolcs Káli

## Abstract

Hippocampal place cells are activated sequentially as an animal explores its environment. These activity sequences are internally recreated (“replayed”), either in the same or reversed order, during bursts of activity (sharp wave-ripples; SWRs) that occur in sleep and awake rest. SWR-associated replay is thought to be critical for the creation and maintenance of long-term memory. In order to identify the cellular and network mechanisms of SWRs and replay, we constructed and simulated a data-driven model of area CA3 of the hippocampus. Our results show that the chain-like structure of recurrent excitatory interactions established during learning not only determines the content of replay, but is essential for the generation of the SWRs as well. We find that bidirectional replay requires the interplay of the experimentally confirmed, temporally symmetric plasticity rule, and cellular adaptation. Our model provides a unifying framework for diverse phenomena involving hippocampal plasticity, representations, and dynamics, and suggests that the structured neural codes induced by learning may have greater influence over cortical network states than previously appreciated.

## 1 Introduction

The hippocampal region plays a pivotal role in spatial and episodic memory (O’Keefe and Nadel, 1978; Morris et al., 1982). The different stages of memory processing (Marr, 1971; Buzsáki, 1989) are associated with distinct brain states, and are characterized by distinct oscillatory patterns of the hippocampal local field potential (LFP) (Buzsáki et al., 1983; Colgin, 2016). When rodents explore their environment, place cells of the hippocampus are activated in a sequence that corresponds to the order in which the animal visits their preferred spatial locations (place fields) (O’Keefe and Dostrovsky, 1971). The same sequences of firing activity can also be identified, on a faster time scale, during individual cycles of the 4-10 Hz theta oscillation that dominates the hippocampal LFP in this state (O’Keefe and Recce, 1993; Dragoi and Buzsáki, 2006; Foster and Wilson, 2007). These compressed sequences are thought to be optimal for learning via activity-dependent synaptic plasticity (Jensen and Lisman, 2005; Foster and Wilson, 2007).

Other behavioral states such as slow-wave sleep and quiet wakefulness are characterized by the repetitive but irregular occurrence of bursts of activity in the hippocampus, marked by the appearance of sharp waves (Buzsáki et al., 1983; Wilson and McNaughton, 1994) and associated high-frequency (ripple) oscillations (O’Keefe and Nadel, 1978; Buzsáki et al., 1992) in the LFP. Disruption of SWRs was shown to impair long-term memory (Girardeau et al., 2009; Ego-Stengel and Wilson, 2010; Jadhav et al., 2012; Oliva et al., 2020). The activity sequences observed during exploration are also recapitulated during SWRs (Nádasdy et al., 1999; Kudrimoti et al., 1999; Lee and Wilson, 2002), in the absence of any apparent external driver, and this “replay” can happen either in the same or in a reversed order relative to the original behaviorally driven sequence (Foster and Wilson, 2006; Csicsvari et al., 2007; Diba and Buzsáki, 2007; Davidson et al., 2009; Karlsson and Frank, 2009; Gupta et al., 2010). More specifically, awake replay is predominantly in the “forward” direction near choice points during navigation (Diba and Buzsáki, 2007; Pfeiffer and Foster, 2013), while it is mainly “backward” when the animal encounters a reward (Diba and Buzsáki, 2007). Consequently, while sleep replay was suggested to be involved in memory consolidation, forward and reverse replay in awake animals may contribute to memory recall and reward-based learning, respectively (Carr et al., 2011; Foster, 2017; Pfeiffer, 2017; Ólafsdóttir et al., 2018). Finally, behavioral tasks with a high memory demand led to an increase in the duration of SWRs, while artificial prolongation of SWRs improved memory (Fernández-Ruiz et al., 2019).

For several decades, area CA3 of the hippocampus has been proposed to be a critical site for hippocampal memory operations, mainly due to the presence of an extensive set of modifiable excitatory recurrent connections that could support the storage and retrieval of activity patterns and thus implement long-term memory (Marr, 1971; McNaughton and Morris, 1987; Levy, 1996; Rolls, 1996; Káli and Dayan, 2000). Area CA3 was also shown to be strongly involved in the generation of SWRs and the associated forward and reverse replay (Buzsáki, 2015; Davoudi and Foster, 2019).

Several modeling studies have addressed either theta oscillogenesis, learning, sequence replay or ripples; however, a unifying model of how the generation of SWRs and the associated neuronal activity are shaped by previous experience is currently lacking. In the present study, we built a minimal, yet data-driven model of the CA3 network, which, after learning via a symmetric spike-timing-dependent synaptic plasticity rule (Mishra et al., 2016) during simulated exploration, featured both forward and backward replay during autonomously generated sharp waves, as well as ripple oscillations generated in the recurrently connected network of perisomatic interneurons. After validating the model against several *in vivo* and *in vitro* results (Buzsáki et al., 1992; Hájos et al., 2013; English et al., 2014; Schlingloff et al., 2014; Stark et al., 2014; Pfeiffer and Foster, 2015; Gan et al., 2017), we took advantage of its *in silico* nature that made feasible a variety of selective manipulations of cellular and network characteristics. Analyzing the changes in the model’s behavior after these perturbations allowed us to establish the link between learning from theta sequences and the emergence of SWRs during “off-line” states, to provide a possible explanation for forward and backward replays, and to disentangle the mechanisms responsible for sharp waves and ripple oscillations.

## 2 Results

To identify the core mechanisms that are responsible for the relationship between learning during exploration and SWRs during rest, we built a scaled-down network model of area CA3 of the rodent hippocampus. In this model, we specifically did not aim to capture all the biological details of hippocampal neurons and circuits; instead, our goal was to reveal and analyze those mechanisms that are essential for the generation of sharp waves, ripple oscillations, and bidirectional activity replay. To gain a mechanistic understanding of the relationship between learning during exploration and SWRs during rest, as well as the generation and role of ripple oscillation, we built a scaled-down network model of area CA3 of the rodent hippocampus. The complete network consisted of 8000 excitatory pyramidal cells (PCs) and 150 inhibitory parvalbumin-containing basket cells (PVBCs), corresponding roughly to the size of the CA3 area in a 600 micrometer-thick slice of the mouse hippocampus, which is known to be capable of generating SWRs (Hájos et al., 2013; Schlingloff et al., 2014). These two cell types are known to be indispensable to the generation of SWRs (Racz et al., 2009; Ellender et al., 2010; English et al., 2014; Schlingloff et al., 2014; Stark et al., 2014; Gulyás and Freund, 2015; Buzsáki, 2015; Gan et al., 2017). All neurons in the network were modeled as single-compartment, adaptive exponential integrate-and-fire (AdExpIF) models whose parameters were fit to reproduce *in vitro* voltage traces of the given cell types in response to a series of step current injections (Methods, Supplementary Figure S2). Connections of all types (PC-PC, PC-PVBC, PVBC-PC, PVBC-PVBC) were established randomly using connection type-specific probabilities estimated from anatomical studies.

Simulations of the network were run in two phases corresponding to spatial exploration and “off-line” hippocampal states (slow-wave sleep and awake immobility), respectively. During the exploration phase, the spiking activity of PCs was explicitly set to mimic the firing patterns of a population of place cells during simulated runs on a linear track, and the recurrent connections between PCs evolved according to an experimentally-constrained, spike-timing-dependent plasticity (STDP) rule (see below). In the subsequent off-line phase, we recorded the spontaneous dynamics of the network in the absence of structured external input (or plasticity), analyzed the global dynamics (including average firing rates and oscillations), and looked for the potential appearance of sequential activity patterns corresponding to the previous activation of place cells during exploration, often referred to as “replay”. The main features of the model are summarized in Figure 1.

**Figure 1:**
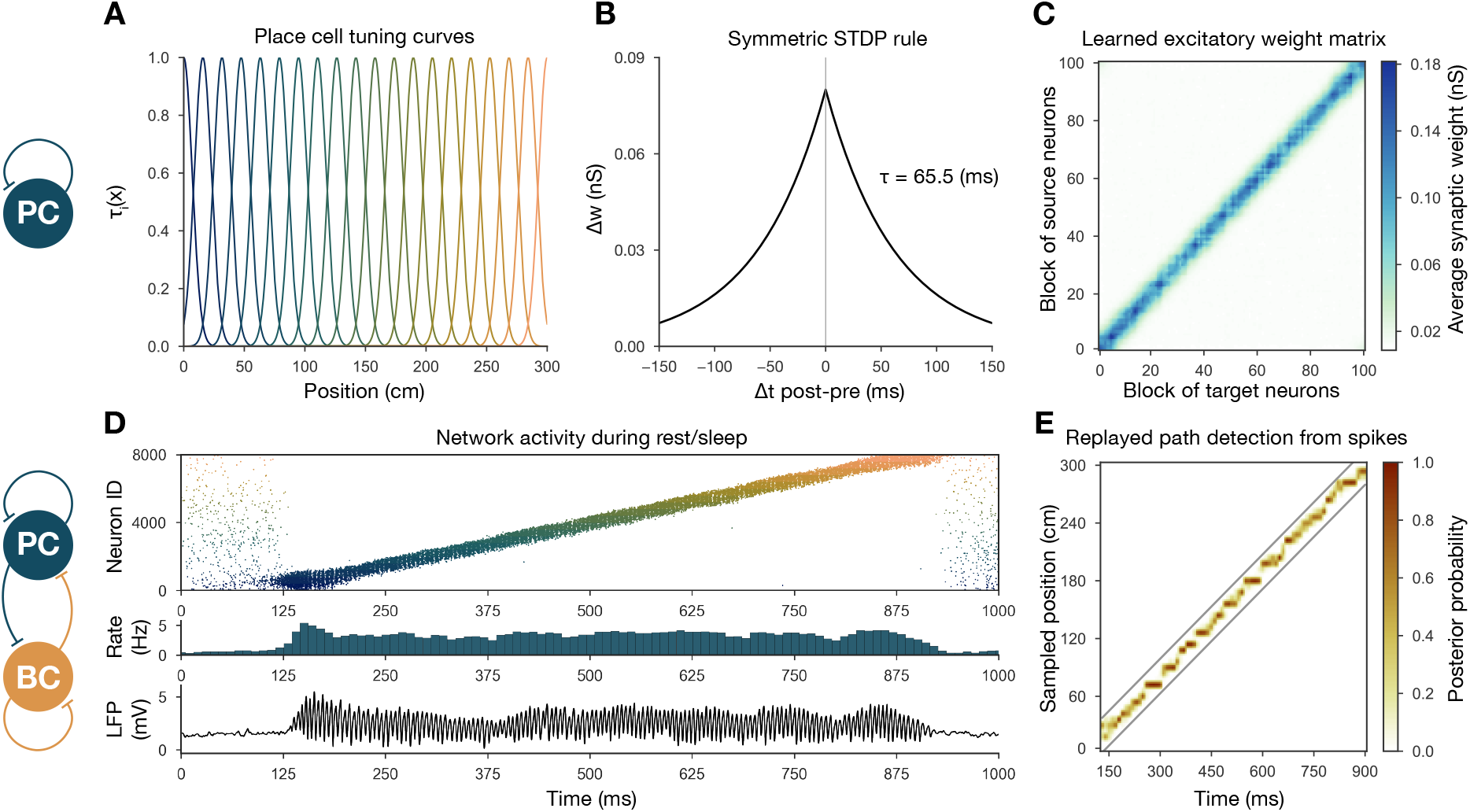
Overview of learning and the spontaneous generation of SWRs and sequence replay in the model. **(A)** Tuning curves (eq. (1)) of exemplar place cells covering the whole 3 m long linear track. **(B)** Broad, symmetric STDP kernel used in the learning phase. The time constant was fit directly to experimental data from Mishra et al. (2016). **(C)** Learned excitatory recurrent weight matrix. Neurons are ordered according to the location of their place fields. Actual dimensions are 8000*8000 (including PCs with no place fields in the environment) but, for better visualization, each pixel shown represents the average of an 80*80 square. **(D)** PC raster plot is shown in the top panel, color-coded and ordered as the place fields in (A). PC population rate (middle), and LFP estimate (bottom panel), corresponding to the same time period. **(E)** Posterior matrix of the decoded positions from spikes within the high activity period shown in (D). Gray lines indicate the edges of the decoded, constant velocity path.

### 2.1 Recurrent weights are learned during exploration via a symmetric STDP rule

During exploration, half of the PCs had randomly assigned, overlapping place fields in the simulated environment, characterized by Gaussian spatial tuning curves, whereas the others fired at low rates in a spatially non-specific manner (Methods, Figure 1A). During simulated unidirectional runs along a 3 meter-long linear track, these tuning curves, modulated by theta oscillation and phase precession, gave rise to generated spike trains similar to those observed for real place cells under similar conditions (Methods, Supplementary Figure S1). The simulated spike trains served as inputs for STDP, which was characterized by the broad, symmetric kernel observed in pairs of CA3 PCs recorded in hippocampal slices (Figure 1B) (Mishra et al., 2016). The anatomical connectivity of PCs was sparse and random, assuming 10% PC-PC connection probability (Lisman, 1999; Andersen et al., 2007), and only these pre-existing connections were allowed to evolve (Methods).

The most prominent feature of the learned recurrent excitatory synaptic weight matrix was its highly organized structure (Figure 1C). Relatively few strong (> 1 nS) synapses near the diagonal (representing pairs of cells with overlapping place fields) emerged from a background of much weaker connections. Similar symmetric weight structures have been used in continuous “bump” attractor models of various neural systems such as head direction cells, place cells, and parametric working memory (Zhang, 1996; Samsonovich and McNaughton, 1997; Káli and Dayan, 2000; Compte et al., 2000). In these studies, the weights were typically either imposed or learned using rate-based (not STDP) learning rules, and led to stationary patterns of activity in the absence of external input. By contrast, previous spiking neuron models of sequence learning used temporally asymmetric STDP rules, resulting in a weight structure dominated by feedforward chains (each neuron giving the strongest input to other neurons which follow it in the spatial sequence), and sequential activity patterns that follow these chains of strong connections (Jahnke et al., 2015; Chenkov et al., 2017). In order to reveal the combined effect of learned, essentially symmetric connections and realistic cell-type-specific neuronal spike responses, we next explored the spontaneously generated activity patterns in our full network model.

### 2.2 The dynamics of the network model reproduce key features of SWRs and replay

The sparse recurrent excitatory weight matrix resulting from the phenomenological exploration (Figure 1C) was used directly in network simulations mimicking resting periods, in which (with the exception of the simulations shown later in Figure 2C) PCs received only spatially and temporally unstructured random synaptic input. The maximal conductances of the other types of connections in the network (the weights of PC-PVBC, PVBC-PC, and PVBC-PVBC connections as well as the weight of the external random input) were optimized using an evolutionary algorithm with network-level objectives, which included physiological PC population firing rates, suppressed gamma oscillation in the PC population and strong ripple oscillation in the PVBC population (see Methods for details).

**Figure 2:**
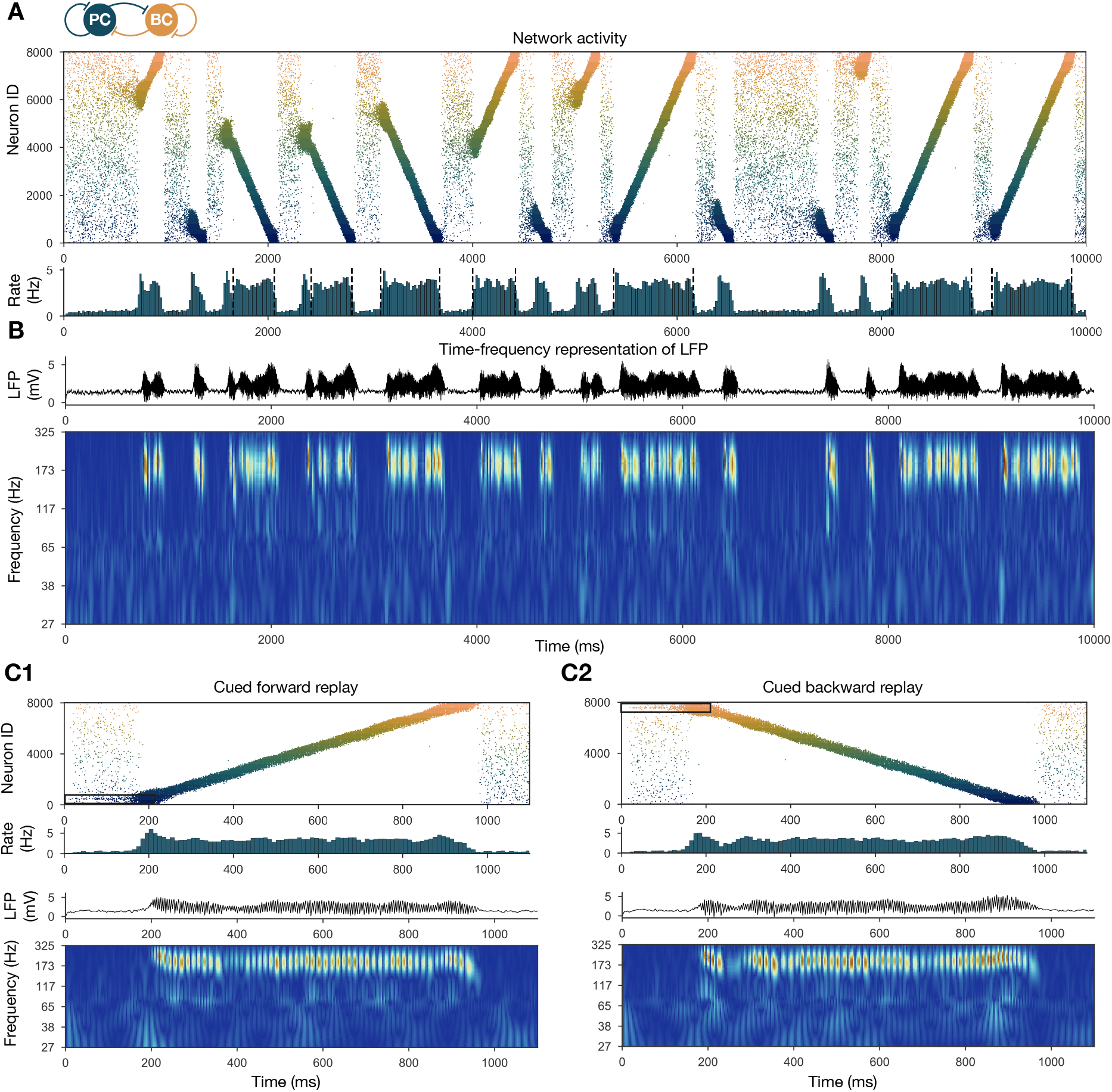
Forward and backward replay events, accompanied by ripple oscillations, can occur spontaneously but can also be cued. **(A)** PC raster plot of a 10-second long simulation, with sequence replays initiating at random time points and positions and propagating either in forward or backward direction on the top panel. PC population firing rate is plotted below. Dashed vertical black lines indicate the periods marked as sustained high activity states (above 2 Hz for at least 260 ms) which are submitted to automated spectral and replay analysis. **(B)** Estimated LFP in the top panel and its time-frequency representation (wavelet analysis) below. **(C)** Forward and backward sequence replays resulting from targeted stimulation (using 200 ms long 20 Hz Poisson spike trains) of selected 100 neuron subgroups, indicated by the black rectangles in the raster plots. **(C1)** Example of cued forward replay. PC raster plot is shown at the top with PC population rate, LFP estimate, and its time-frequency representation (wavelet analysis) below. **(C2)** Same as (C1), but different neurons are stimulated at the beginning, leading to backward replay.

Spontaneously generated activity in the network consisted of alternating periods of low activity (mean PC rates below 1 Hz) and high activity (mean PC rates around 3.5 Hz), which resembled the recurring appearance of sharp wave events during slow-wave sleep and quiet wakefulness (Figure 2A). Similar to experimental sharp waves, high-activity events in the model were accompanied by transient oscillations in the ripple frequency range (Figure 2B) (Buzsáki, 1986; Buzsáki et al., 1992; Foster and Wilson, 2006; Buzsáki, 2015). We calculated an estimate of the LFP by summing the synaptic inputs of a small, randomly selected subset of PCs (Mazzoni et al., 2008), and found that ripple oscillations were reliably present in this signal during the high-activity periods (Figure 1D, Figure 2B).

When we plotted the spike times of all PCs with place cells ordered according to the location of their place fields on the track during the learning phase, it became clear that place cell sequences were replayed during simulated SWRs, while activity had no obvious structure in the low-activity periods between SWRs (Figure 2A). Interestingly, replay of place cell sequences could occur in either the forward or the backward direction relative to the order of activation in the learning phase (Figure 2A). These qualitative observations were confirmed by analysing sequence replays with a Bayesian place decoding and path fitting method, using spatial tuning curves introduced for spike train generation in the exploration part (Methods, Figure 1E) (Davidson et al., 2009; Karlsson and Frank, 2009; Ólafsdóttir et al., 2018). Interestingly, our network model also reproduced the broad, long-tailed step size distribution in the sequence of decoded locations during SWRs, as observed experimentally by Pfeiffer and Foster (2015) (Supplementary Figure S3).

It was also possible to “cue” replay, by giving an additional external stimulation to a small subpopulation (*n* = 100) of PCs which had overlapping place fields in the learned environment. In this case, sequence replay started at the location corresponding to the cells which received extra stimulation (Figure 2C). This feature may explain why awake replay tends to be forward when the animal is planning a trajectory from its current location, and backward at the goal location.

At the level of single cells, we found that both PCs and PVBCs received approximately balanced excitation and inhibition during SWR events, and inhibitory currents were modulated at the ripple frequency in these periods (Supplementary Figure S4C). Excitatory inputs dominated between SWRs, and during the initiation of SWR events. Only a small minority of PCs fired in individual ripple cycles, while the participation of PVBCs was much higher (but not complete), resulting in a mean PVBC firing rate of ~65 Hz during the SWRs, which is much higher than their baseline rate, but significantly below the ripple frequency (Supplementary Figure S4A, B). All of these findings were consistent with experimental results *in vitro* (Hájos et al., 2013; Schlingloff et al., 2014) and *in vivo* (English et al., 2014; Hulse et al., 2016; Gan et al., 2017). On the other hand, we note that the single cell firing rate distribution of place cells was close to normal in the model (Supplementary Figure S4A1), different from the lognormal distribution reported *in vivo* (Mizuseki and Buzsáki, 2013).

#### 2.2.1 SWRs and replay are robust when recurrent excitation is varied

To show that our network model reproduces the wide range of experimental findings presented above in a robust manner, we ran a sensitivity analysis of several parameters. We started by studying the effects of the recurrent excitatory weights, which are the link between exploratory and resting dynamics. To this end, we ran simulations with systematically up- and downscaled PC-PC weights, and automatically evaluated various features of the network dynamics, such as population-averaged firing rates, the presence of sequence replay, as well as significant peaks in the ripple frequency range in the power spectra as shown before (Methods, Figure 3). The network with PC-PC weights multiplied by 0.8 displayed a low-activity, noisy state with severely reduced mean PC firing rate, no sequence replay, and no clear oscillation (Figure 3A1). At the 0.9 multiplier level sequence replays started to appear, but were less frequent than in the baseline model (Figure 3A2). As the PC-PC synaptic weights were scaled up, sequence replays became faster (completed in a shorter time) and occurred more often (Figure 3A3), a behavior which was stable and realistic up to the 1.5 multiplier level. Ripple oscillations had higher power and appeared at lower multiplier levels in the PVBC population than in the PC population (Supplementary Figure S5), suggesting that they originate from the PVBCs and propagate to the PCs, in agreement with theories based on experimental findings (Buzsáki et al., 1992; Ylinen et al., 1995; Racz et al., 2009; Schlingloff et al., 2014; Stark et al., 2014; Gan et al., 2017).

**Figure 3:**
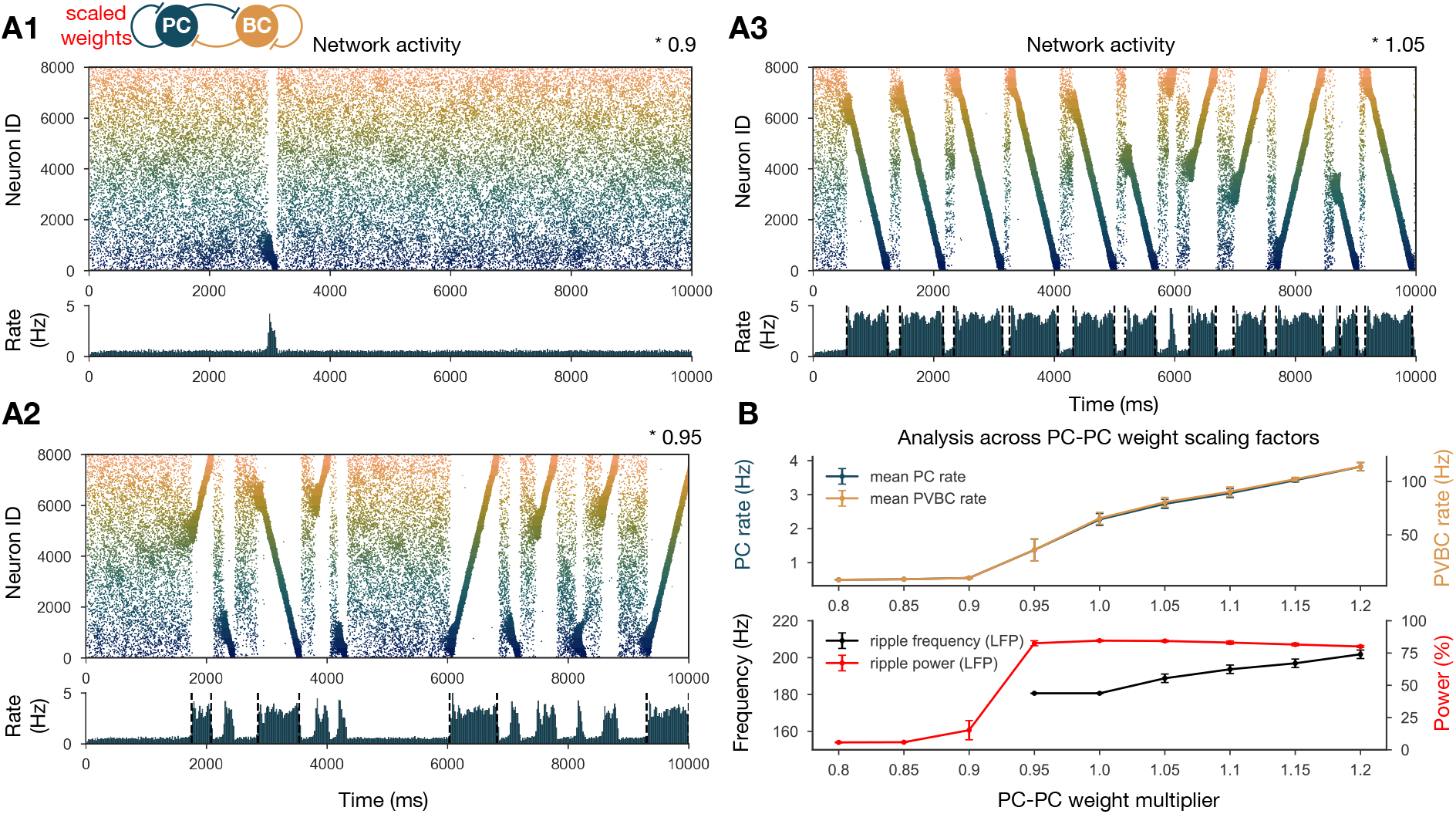
Sharp waves, ripple oscillations, and replay are robust with respect to scaling the recurrent excitatory weights. **(A)** PC raster plots on top and PC population rates at the bottom for E-E scaling factors: 0.9 (A1), 0.95 (A2) and 1.05 (A3). (Scaling factor of 1.0 is equivalent to Figure 2A.) Dashed vertical black lines have the same meaning as in Figure 2A. **(B)** Analysis of selected indicators of network dynamics across different E-E weight scaling factors (0.8-1.2). Mean PC (blue) and PVBC (gold) population rates are shown on top. The frequency of significant ripple oscillations (black) and the percentage of power in the ripple frequency range (red) in the estimated LFP is shown at the bottom. Errors bars indicate standard deviation and are derived from simulations with 5 different random seeds. See also Supplementary Figure S5.

#### 2.2.2 Multiple environments can be learned and replayed

Next we showed that it is possible to store the representations of multiple environments in the weight matrix, and this change does not fundamentally alter the network dynamics (Figure 4). In particular, we simulated experience and STDP-based learning in two linear environments with a different but overlapping random set of place cells (Figure 4A). The resulting population-level dynamics was quite similar to the one following experience in a single environment, but, during each SWR event, a place cell sequence from either one or the other environment was reactivated (Figure 4B, C). Learning two sequences with the same additive symmetric (non-decreasing) STDP rule led to stronger PC-PC synapses on average (Figure 4A4), which resulted in a higher overall mean PC rate (Figure 4B, D). As a consequence, detectable sequence replays and significant ripple oscillations appeared at lower PC-PC weight multiplier levels (Figure 4D).

**Figure 4:**
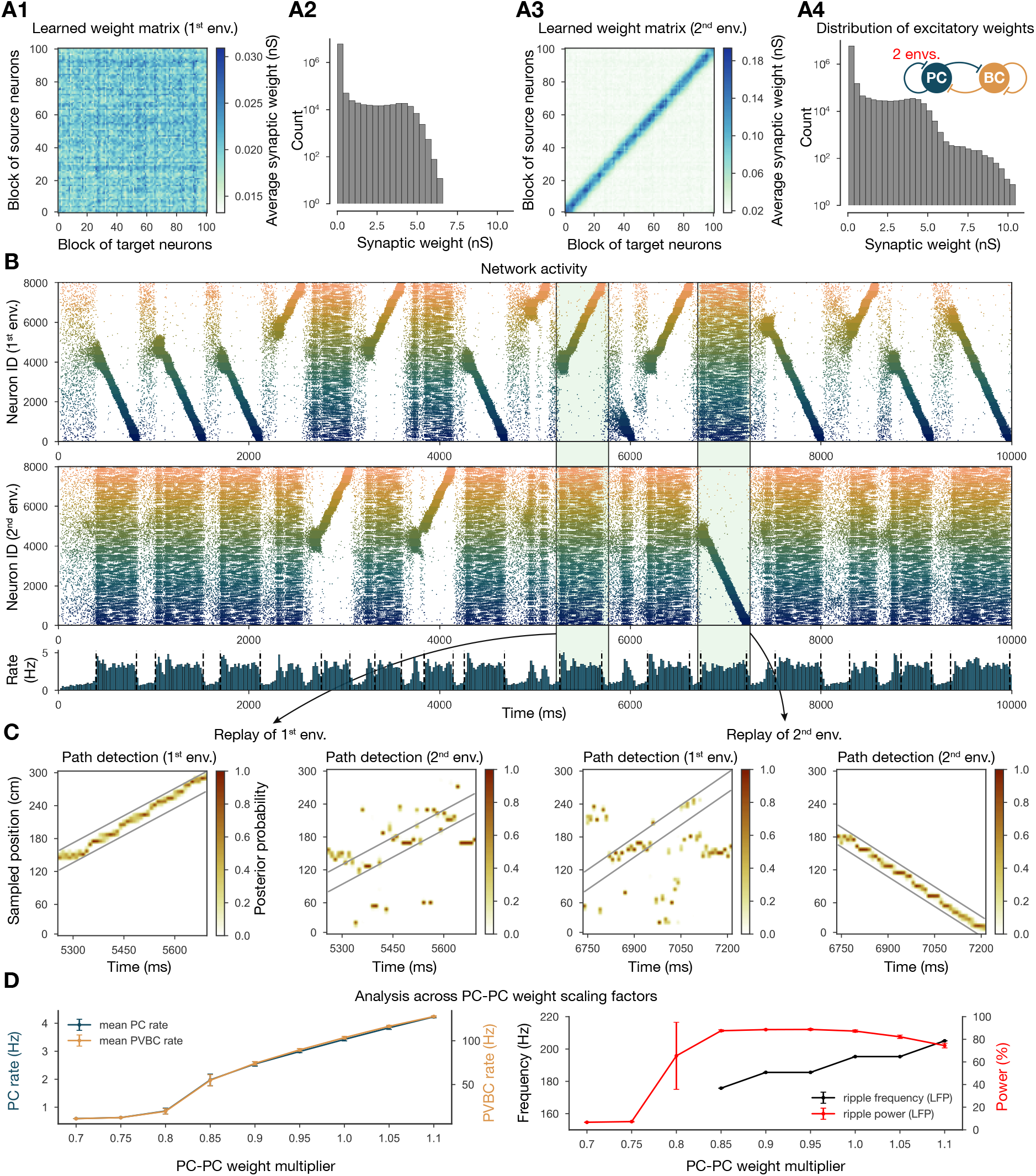
Two distinct environments can be learned and replayed by the network. **(A)** Learned excitatory recurrent weight matrices. (A1) Weights after learning the first environment. Note that the matrix appears random because neurons are arranged according to their place field location in the second environment, which has not been explored at this point. (A3) Weights after learning in the second environment. (A2) and (A4) distribution of non-zero synaptic weights in the learned weight matrices in (A1) and (A2) respectively. **(B)** PC raster plots: in the top panel neurons are ordered and colored according to the first environment; in the middle panel neurons are ordered and colored according to the second environment; and PC population rate is shown at the bottom (see Figure 2A) from a simulation run with 0.9* the modified weight matrix shown in (A3). **(C)** Posterior matrices of decoded positions from spikes (see Figure 1E) within a selected high activity state (8th and 10th from (B)). From left to right: decoding of replay in 1st environment (8th event from (B)) according to the first (significant) and second environment; decoding of replay in second environment (10th event from (B)) according to the first and second (significant) environment. **(D)** Analysis of selected network dynamics indicators across different E-E weight scaling factors (0.7-1.1) as in Figure 3 (B).

### 2.3 Manipulating the model reveals mechanisms of sharp wave-ripple generation and sequence replay

#### 2.3.1 Symmetric STDP rule is necessary for bidirectional replay

Our network model, equipped with structured recurrent excitation resulting from learning, was able to robustly reproduce recurring sharp wave events accompanied by bidirectional sequence replay and ripple oscillations. Next, we set out to modify this excitatory weight matrix to gain a more causal understanding of how the learned weight pattern is related to the emergent spontaneous dynamics of the network.

First, in order to gauge the significance of the experimentally determined symmetric STDP rule, we changed the STDP kernel to the classical asymmetric one that characterizes many other connections in the nervous system (Figure 5A-C) (Bi and Poo, 1998; Gerstner et al., 2014). In this case, the learned weight matrix was reminiscent of the feedforward chains that characterized several of the earlier models of hippocampal replay (Jahnke et al., 2015; Chenkov et al., 2017; Theodoni et al., 2018). We found that this weight structure also supported the generation of SWR events and the reactivation of learned sequences in our model; however, crucially, sequence replays occurred only in the forward direction (Figure 5D, E). Theodoni et al. (2018) presented a thorough analysis of the relationship between the shape of the plasticity kernel and the sequence replay in a rate based model, and thus we shifted our focus towards different modifications.

**Figure 5:**
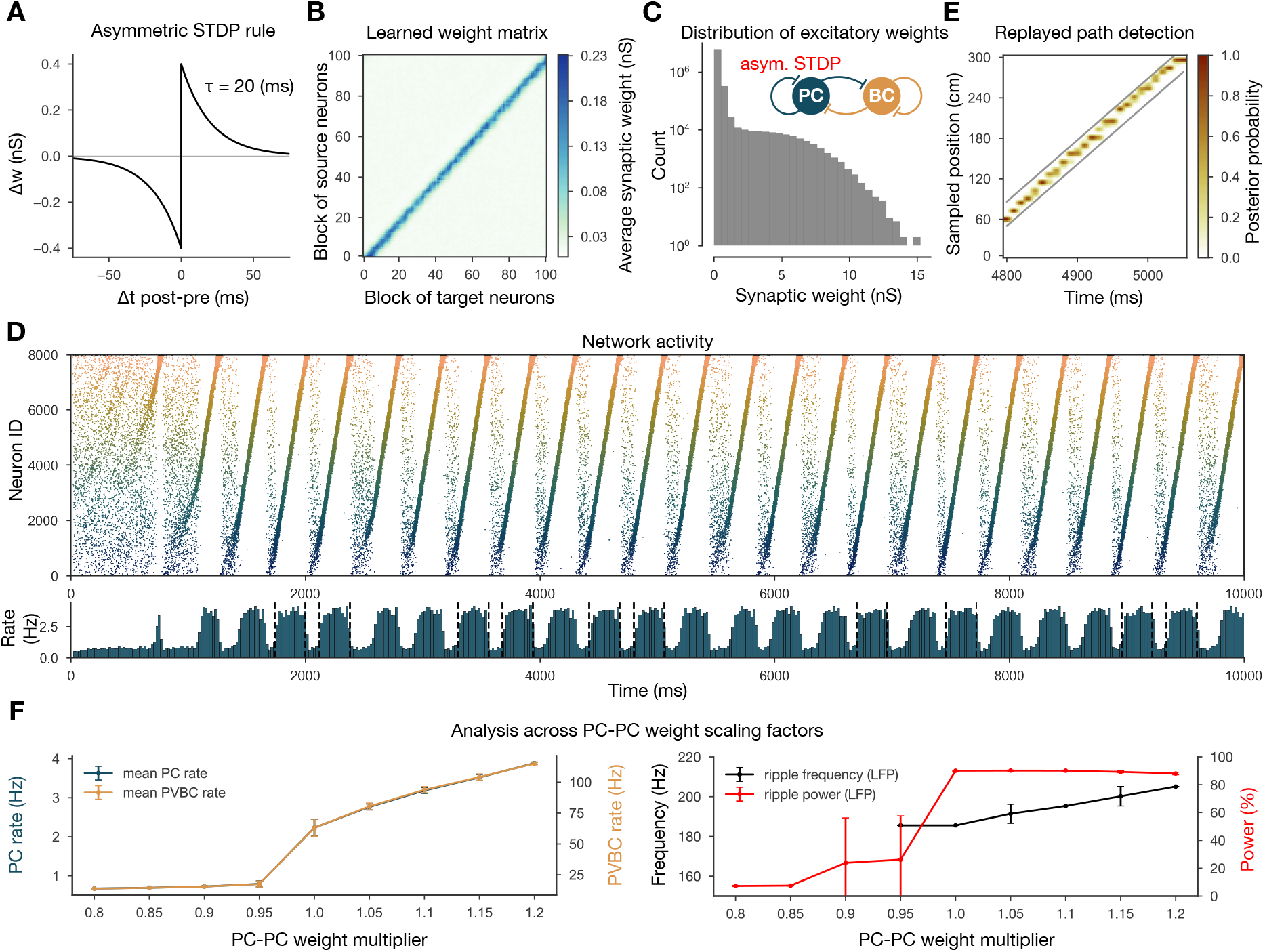
Learning with an asymmetric STDP rule leads to the absence of backward replay. **(A)** Asymmetric STDP kernel used in the learning phase. **(B)** Learned excitatory recurrent weight matrix. **(C)** Distribution of non-zero synaptic weights in the weight matrix shown in (B). **(D)** PC raster plot on top and PC population rate at the bottom (see Figure 2 (A) from a simulation run with the weight matrix shown in (B). **(E)** Posterior matrix of the decoded positions from spikes (see Figure 1 (E) within a selected high activity state (6th one from (D)). **(F)** Analysis of selected network dynamics indicators across different E-E weight scaling factors (0.8-1.2) as in Figure 3 (B).

#### 2.3.2 The structure rather than the statistics of recurrent excitatory weights is critical for SWRs

Inspired by the observation that many network-level properties, such as single PC firing rates, burst index, and participation in SWR events, follow a skewed, lognormal distribution *in vivo* (Mizuseki and Buzsáki, 2013), Omura et al. (2015) built a network model with recurrent excitatory weights following a lognormal distribution. Their network with unstructured, but lognormally distributed recurrent synaptic strengths reproduced most of the *in vivo* observations of Mizuseki and Buzsáki (2013); however, no sequence replay or ripple oscillation was involved. In the network presented here, the distribution of PC-PC weights is the result of the application of STDP on the generated spike trains and does not strictly follow a lognormal distribution, although it has a similar long tail (Figure 6D2). In order to establish whether the overall distribution or the fine structure of the weights is the key determinant of neural dynamics in our model, we performed two more drastic perturbations of the recurrent weight matrix itself, starting from the version established in the learning phase.

**Figure 6:**
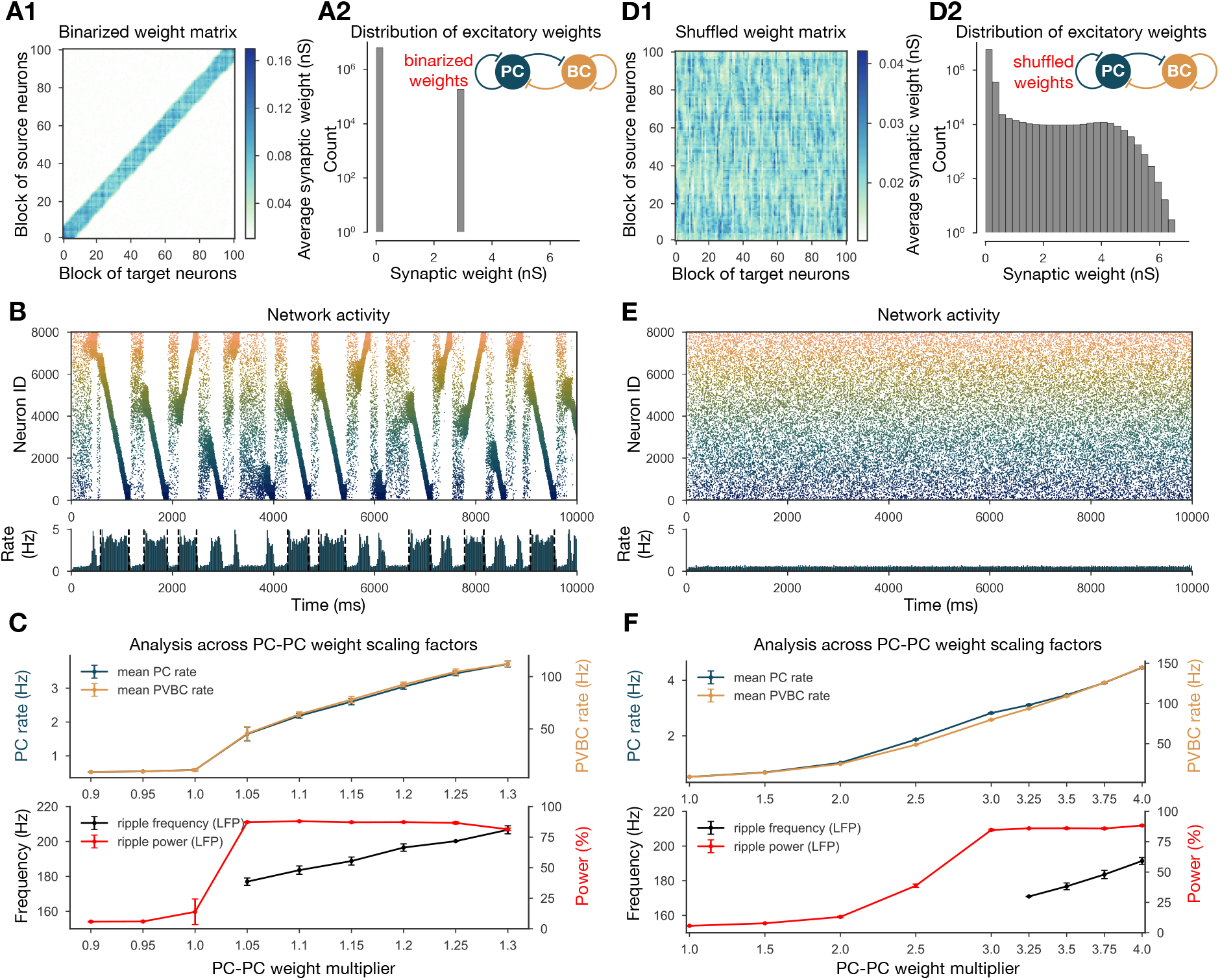
Altering the structure of recurrent excitatory interactions changes the network dynamics but altering the weight statistics has little effect. **(A1)** Binarized (largest 3% and remaining 97% non-zero weights averaged separately) recurrent excitatory weight matrix. (Derived from the baseline one shown in Figure 1C.) (A2) Distribution of non-zero synaptic weights in the learned weight matrix shown in (A1). **(B)** PC raster plot on top and PC population rate at the bottom (see Figure 2A) from a simulation ran with 1.1* the binarized weight matrix shown in (A). **(C)** Analysis of selected network dynamics indicators across different E-E weight scaling factors (0.9-1.3) as in Figure 3B. **(D1)** Column-shuffled recurrent excitatory weight matrix. (Derived from the baseline one shown in Figure 1C.) **(D2)** Distribution of non-zero synaptic weights in the weight matrix shown in (D1) (identical to the distribution of the baseline weight matrix shown in Figure 1C). **(E)** PC raster plot on top and PC population rate at the bottom from a simulation run with the shuffled weight matrix shown in (D1). **(F)** Analysis of selected network dynamics indicators across different E-E weight scaling factors (1.0-4.0) as in Figure 3B. Note the significantly extended horizontal scale compared to other cases.

Our first perturbation kept the structure of the interactions and the overall mean weight intact, but completely changed the distribution of the weights. This was achieved by binarizing the values of the learned weight matrix. Specifically, we divided weights into two groups, the strongest 3% and the remaining 97%, and set each weight in both groups to the group average (Figure 6A). Using the modified recurrent synapses between the PCs in the network, we observed extremely similar behaviour to our baseline network: sequence replays in both directions, always accompanied by ripple oscillation, with only a small change in the required multiplier for the PC-PC weights (Figure 6B, C).

The second modification kept the same overall weight distribution and even the actual values of the outgoing weights for all neurons, but destroyed the global structure of synaptic interactions in the network. To this end, we randomly shuffled the identity of the postsynaptic neurons (by shuffling the columns of the weight matrix). Strong synapses were not clustered anymore along the diagonal (representing interactions between neurons with nearby place fields), but distributed homogeneously within the matrix (Figure 6D1). None of the networks equipped with the scaled versions of this shuffled weight matrix exhibited sequence replay, mean PC rates were severely reduced, and no sharp wave-like events were observed (Figure 6E, F). On the other hand, with sufficiently amplified (*3.5) PC-PC weights we detected significant peaks in the ripple frequency range of the LFP (Figure 6F).

Taken together these modifications suggest that, unlike in the model of Omura et al. (2015), the distribution of the excitatory recurrent synaptic weights is neither necessary nor sufficient for the observed physiological population activity in our model. In other words, our simulation results suggest that the fine structure of recurrent excitation not only enables coding (sequence replay), but also has a major effect on the global average network dynamics (firing rates, sharp waves and ripple oscillations) in hippocampal area CA3.

#### 2.3.3 Cellular adaptation is necessary for replay

As mentioned earlier, most previous models with symmetrical local excitatory interactions and global feedback inhibition functioned as “bump attractor” networks, in which the dynamics converge to stable patterns of activity involving high rates in a group of neurons with similar tuning properties (adjacent place fields), and suppressed firing in the rest of the excitatory population (Zhang, 1996; Samsonovich and McNaughton, 1997; Káli and Dayan, 2000; Compte et al., 2000). Recent work with rate-based models has also shown that these “stable bumps” can be transformed into “traveling bumps” by the introduction of short-term depression (York and van Rossum, 2009; Romani and Tsodyks, 2015; Theodoni et al., 2018) or spike threshold adaptation (Itskov et al., 2011; Azizi et al., 2013). On the other hand, previous spiking models of sequence learning and recall/replay typically relied on temporally asymmetric learning rules and the resulting asymmetric weight matrices to ensure that neurons are reactivated in the same sequence as during learning (Jahnke et al., 2015; Chenkov et al., 2017), which is also why it is difficult for these models to capture bidirectional replay. By contrast, our model uses the experimentally recorded symmetric STDP rule, which results in symmetrical synaptic interactions (although only at the population level rather than the single neuron level, due to the randomness of connectivity). Since our network generated a bump of activity that traveled unidirectionally in any given replay event rather than a “stable bump”, we hypothesized that the cellular-level adaptation that characterized CA3 PCs and was also captured by our model may destabilize stable bumps and lead to their constant movement.

To test this hypothesis, we re-fitted our single-cell data on PC responses to current injections using a modified ExpIF model which did not contain any adaptation mechanism (but was otherwise similar to the baseline model). Although the non-adapting ExpIF model provided a reasonably good fit to our single-cell data (Figure 7A), and the weights resulting from learning were identical to the baseline case, the spontaneous network dynamics was completely different: there was no sequence replay for any scaling of the recurrent PC-PC weights; instead, when structured activity emerged from the random background, it was in the form of a stationary rather than a moving bump (Figure 7B). Therefore, it was the combination of a symmetric learning rule with cellular adaptation that created the possibility of bidirectional replay in our network model.

**Figure 7:**
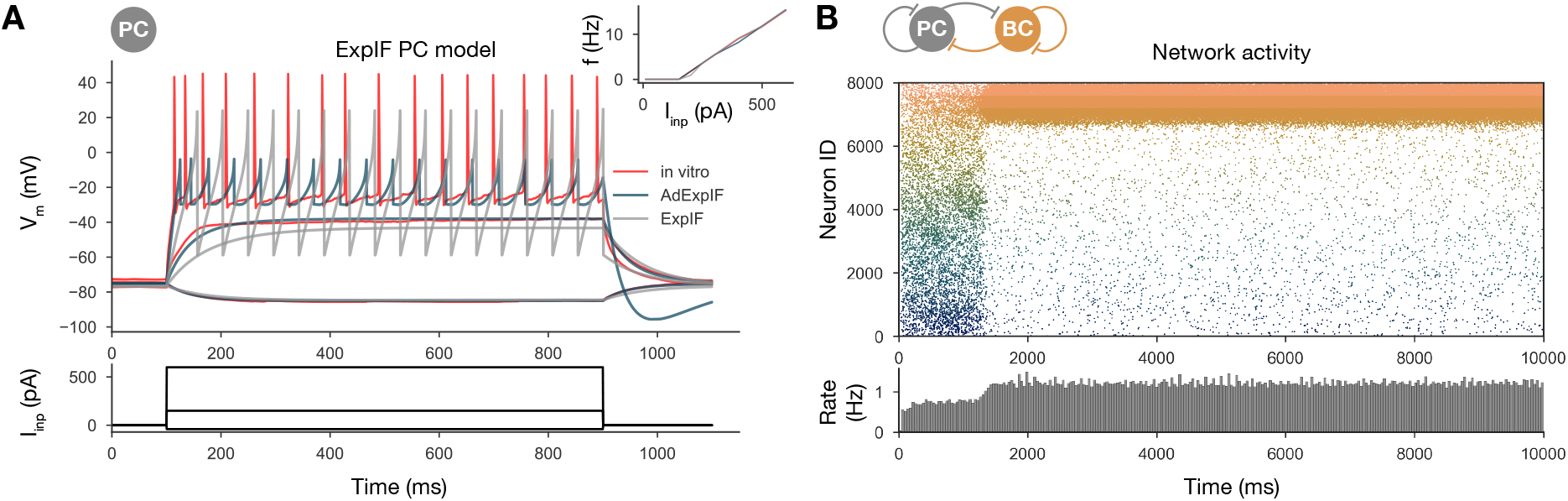
Sequential replay requires firing rate adaptation in the PC population. **(A)** Voltage traces of fitted AdExpIF (blue) and ExpIF (gray) PC models and experimental traces (red) are shown in the top panel. Inserts show the fI curves of the *in vitro* (red) and *in silico* cells. The amplitudes of the 800 ms long step current injections shown at the bottom were as follows: −0.04, 0.15 and 0.6 nA. For parameters of the cell models see Table 2. **(B)** PC raster plot of a 10-second long simulation with the ExpIF PC models, showing stationary activity in the top panel. PC population rate is shown below.

#### 2.3.4 Ripple oscillations are generated by the recurrently coupled inhibitory population

From the weight matrix modifications, we also learned that ripple oscillations can be disentangled from sequence replays, and only require sufficient drive to the interconnected PVBC population in our model (Figure 6F). The same conclusion was reached by recent experimental studies *in vivo* (Stark et al., 2014) and *in vitro* (Ellender et al., 2010; Schlingloff et al., 2014). To further investigate the generation of ripples in our model, we simulated and analyzed two additional modified versions of the full network.

First, we disconnected the PVBC network from the PCs and replaced the PC input with independent Poisson synaptic input at rates derived from the firing of the PC population during SWRs in the full simulation (Figure 8A, B). In this simplified, recurrently connected, purely PVBC network we systematically scanned different values of PC input rate and PC-PVBC synaptic weight and measured ripple power as well as the frequency of any significant ripple oscillation as in the full network before (Figure 8A, B). We found that ripple oscillations emerged when the net excitatory drive to PVBCs (which is proportional to the product of the incoming weight and the presynaptic firing rate) was sufficiently large. Above this threshold, the properties of the oscillation depended only mildly on the strength of the input (e.g., the frequency of the oscillation increased moderately with increasing drive), and the firing of PVBCs was synchronized mainly by the decay of inhibitory synaptic current evoked by shared input from their peers.

**Figure 8:**
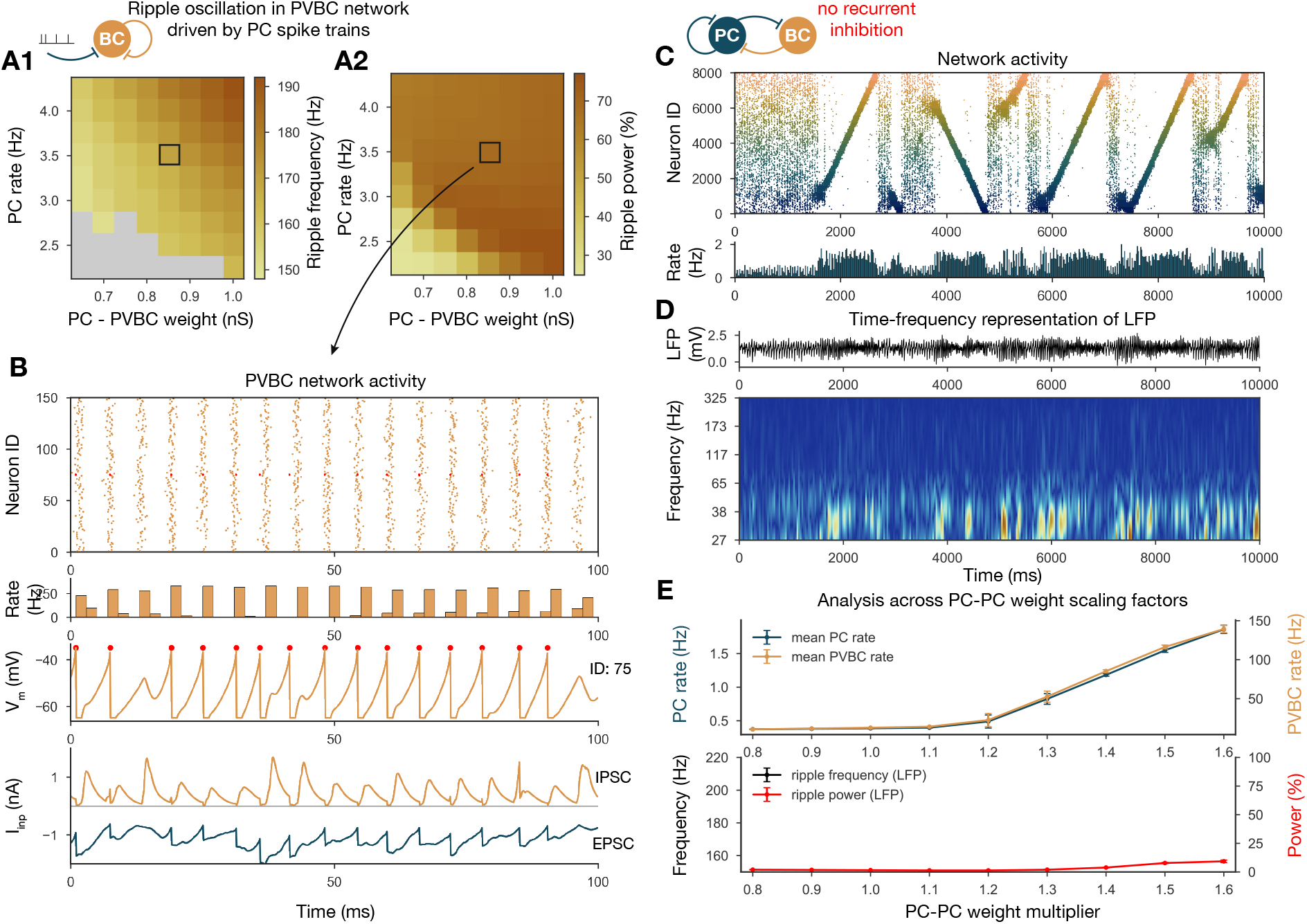
Generation of ripple oscillations relies on recurrent connections within the PVBC population. **(A)** Significant ripple frequency (A1) and ripple power (A2) of a purely PVBC network, driven by (independent) spike trains mimicking PC population activity. Gray color in (A1) means no significant ripple peak. **(B)** From top to bottom: Raster plot, mean PVBC rate, voltage trace (of a selected cell), EPSCs and IPSCs of the selected cell from the middle (100 ms long window) of a simulation used for (A). Ripple frequency and power corresponding to this simulation are marked with a black rectangle in (A1) and (A2). **(C)** PC raster plot on top and PC population rate at the bottom from a simulation ran with a network with 1.3* baseline E-E weight matrix but without any PVBC-PVBC synapses featuring stochastic forward and backward replays but no ripple oscillation (see below). **(D)** Estimated LFP in the top panel and its time-frequency representation (wavelet analysis) below. (Compared to Figure 2B there is increased power in the gamma range, but no ripple frequency oscillation.) **(E)** Analysis of selected network dynamics indicators across different E-E weight scaling factors (0.8-1.6) as in Figure 3B.

Several experiments have shown that local application of GABA blockers eliminates ripples (Maier et al., 2003; Ellender et al., 2010; Schlingloff et al., 2014; Stark et al., 2014); however, in an experimental setup it is hard to distinguish feedback inhibition (PVBC-PC) from reciprocal inhibition (PVBC-PVBC). As a final perturbation, we modified the full baseline model by eliminating only the recurrent inhibitory synapses (Figure 8C, D). The resulting dynamics were stable and with enhanced (*1.3) PC-PC weights it also displayed sequence replay, but ripple oscillations were never observed (Figure 8D, E). Taken together, these results support the conclusions of previous modeling (Brunel and Wang, 2003; Geisler et al., 2005; Taxidis et al., 2012; Donoso et al., 2018; Ramirez-Villegas et al., 2018) as well as experimental studies (Buzsáki et al., 1992; Ylinen et al., 1995; Racz et al., 2009; Ellender et al., 2010; Schlingloff et al., 2014; Stark et al., 2014; Gulyás and Freund, 2015; Gan et al., 2017) proposing that ripple oscillations are generated in strongly driven, recurrently connected inhibitory networks by the fast inhibitory neuronal oscillation (FINO) mechanism. In fact, recurrent inhibitory connections were both necessary and sufficient for the generation of ripple oscillations in our model.

## 3 Discussion

Using a data-driven network model of area CA3 of the hippocampus which reproduces the main characteristics of SWRs, we examined the link between learning during exploration and the network dynamics in resting periods. Our principal findings from analyzing and manipulating this model are as follows: (1) structured (learned) recurrent excitation in the CA3 region not only enables coding and memory, but is critical for the generation of SWRs as well; (2) the symmetric STDP rule described by Mishra et al. (2016), in combination with cellular adaptation in CA3 PCs, provides an explanation for the coexistence of forward and reverse replays; (3) the pattern of strong connections in the network rather than the overall weight statistics may be critical for the emergence and key properties of SWRs and replay in area CA3; (4) ripple oscillations are generated in the strongly driven, recurrently connected network of fast-spiking PVBCs by the FINO mechanism (Schlingloff et al., 2014) (also known as PYR-INT-INT (Stark et al., 2014; Buzsáki, 2015; Ramirez-Villegas et al., 2018)).

### 3.1 Connections of sharp waves, sequence replay and ripple oscillations

SWRs represent the most synchronous physiological activity pattern with the largest excitatory gain in the mammalian brain (Buzsáki et al., 1983, 1992; Buzsáki, 1989, 2015). Under normal conditions, ripples are typically observed riding on top of naturally emerging sharp waves. More recently, using optogenetics, Schlingloff et al. (2014) and Stark et al. (2014) managed to decouple ripples from sharp waves by directly activating the interconnected network of PVBCs. Our *in silico* results perfectly parallel this work: without drastic, non-physiological modifications of the model ripples were always tied to sequence replay, which was in turn associated with bouts of increased spiking activity in the network (the sharp waves). When we separated the BC network, we found that a relatively high (> 2 Hz) mean firing rate in the PC population was required for inducing ripple oscillation, a condition that was satisfied only during sharp wave events and the associated sequence replay in our full baseline network. When the PC population reaches this frequency after a stochastically initiated buildup period, the strongly driven, high-frequency firing of PVBCs is synchronized and phase-locked via reciprocal inhibition. Thus, the learned recurrent PC-PC synaptic weights are responsible for coding, govern sequence replay and, by giving rise to high PC activity during the replay, they also cause the ripples. In summary, memory storage and recall, as well as the main hippocampal oscillations and transient activity patterns, are intimately interconnected in our unifying model.

### 3.2 Biological plausibility of the model

The network model presented here was constrained strongly by the available experimental data. Many cellular and synaptic parameters were fit directly to *in vitro* measurements, and most functional parameters correspond to *in vivo* recordings of hippocampal place cells. Nevertheless, there are certainly many biological features that are currently missing from our model. We do not see this as a major limitation of our study, as our goal was to provide a mechanistic explanation for a core set of phenomena by identifying the key underlying biological components and interactions. On the other hand, our model can be systematically refined and extended, especially when new experimental data become available. The main assumptions we made when constructing the model are explicitly stated in Table 1. Here we briefly discuss some of these assumptions, as well as some remaining discrepancies between our simulation results and the corresponding experimental data.

**Table 1:**
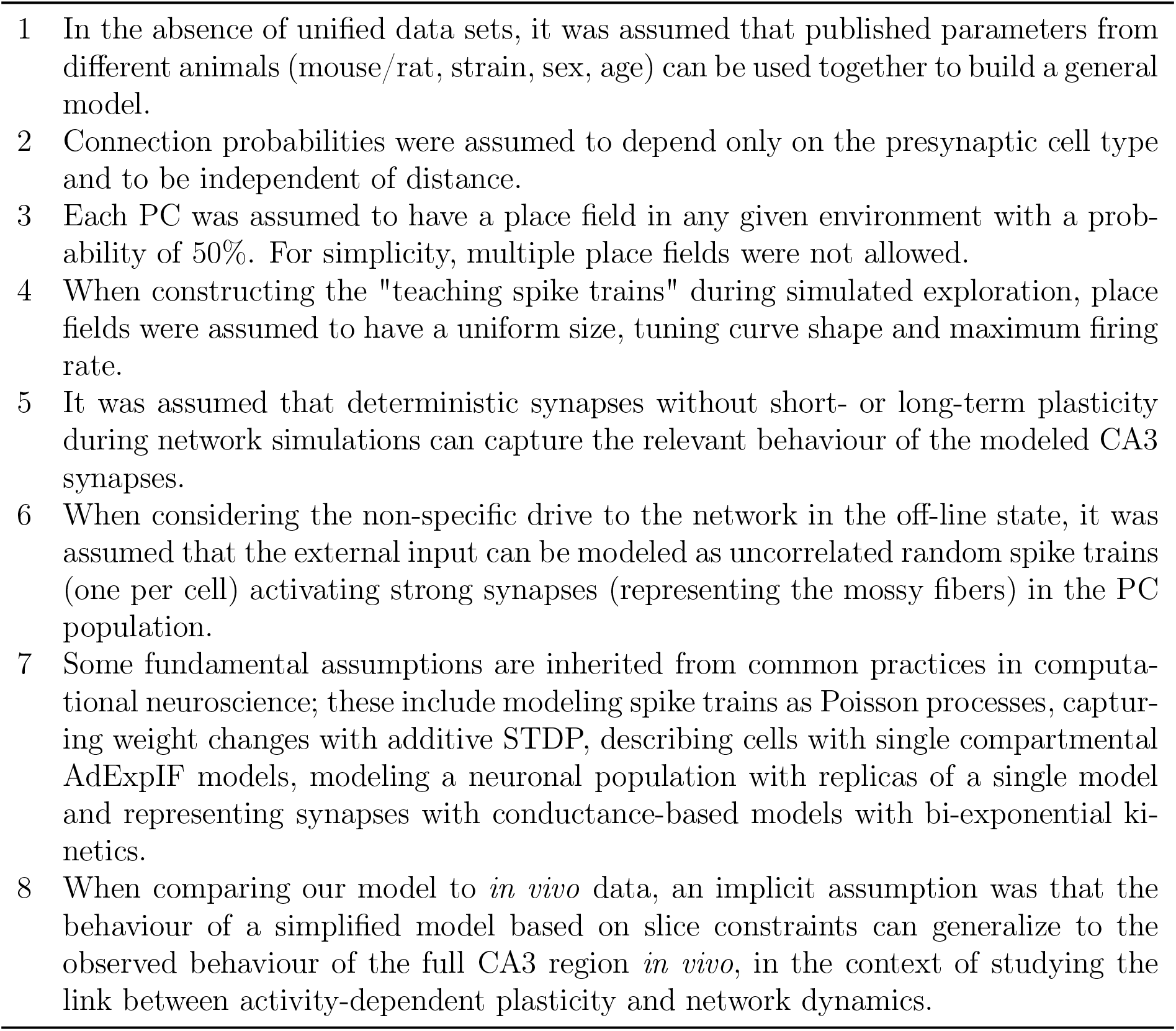
List of modeling assumptions.

The 10% PC-PC connection probability is based on the classical viewpoint that considers the CA3 region as a highly interconnected network (Lisman, 1999; Andersen et al., 2007). Although a recent physiological study Guzman et al. (2016) estimated < 1% connection probability in 400 *μm* thick slices, the authors concluded from “virtual slicing” experiments that recurrent axons were substantially reduced. This is in agreement with Li et al. (1994), who reported at least 70% axonal loss in 400 *μm* slices. Thus, the *in vivo* connection probability is likely to be considerably higher than 1%.

Our model network contained 8000 PCs and 150 PVBCs, which is rather small compared to the full rodent CA3 region. While *in vitro* studies suggest that this network size is sufficient for the generation of SWRs, and that these two cell types are the key players in this process, it is likely that the much larger network size and the various additional cell types modify the quantitative aspects of SWRs *in vivo*.

For simplicity, we also decided not to model all the different types of variation in single cell and synaptic properties that characterize real cortical networks. However, it is important to note that several sources of variability (e.g., those resulting from the random assignment of place field locations, sparse, randomized connectivity, and stochastic, independent external drive) were still included in the model. The variability that we include is sufficient to avoid the regular, unnaturally synchronous dynamics that often characterize homogeneous networks, and results in variable, natural-looking spike patterns during both exploration and spontaneous SWR activity. Including additional forms of heterogeneity would likely increase the cell-to-cell variability of activity (such as firing rates), but (up to a certain limit) would not be expected to change our results qualitatively.

We modeled activity-dependent synaptic plasticity based on the study of Mishra et al. (2016), who uncovered an unusual, temporally symmetric form of spike-timing-dependent plasticity in the recurrent excitatory connections between CA3 pyramidal neurons. We fit the temporal kernel of our STDP rule directly to their data, assumed linear summation of synaptic modifications from all the spike pairs that occurred during the exploration phase, and ignored all higher-order (triplet, etc.) spike interactions. However, we note that the details of the plasticity rule are not expected to have a major impact on our results concerning the emergent dynamics of the CA3 network during resting periods, as long as learning results in the strengthening of connections between PCs that are approximately co-active during exploration, independent of the exact temporal order of their spikes. This requirement would be satisfied by a variety of different learning rules, including the behavioral time-scale plasticity rule recently described in hippocampal CA1 PCs (Bittner et al., 2017).

One substantial difference between SWRs in the model and those recorded *in vivo* is the duration of the SWR events and the associated replay. Ripple episodes typically last 40-100 ms *in vivo* (O’Keefe and Nadel, 1978; Buzsáki et al., 1983, 1992; Ylinen et al., 1995), although Fernández-Ruiz et al. (2019) recently showed that learning in spatial memory tasks is associated with prolonged SWRs and replays. In our model, SWRs can be up to 800 ms in duration as they are terminated when the replayed sequence reaches either the end or the beginning of the learned trajectory (depending on the direction of replay), thus the length of the track determines the maximal duration of the SWR, in combination with the speed of replay (i.e., the rate at which activation propagates across the population of place cells). The speed of replay in the model is consistent with experimental findings, but a single replay event can cover the whole 3m-long track. Davidson et al. (2009) also used a long track in their experiments; however, they reported that the whole path was never replayed during a single SWR, only short segments, which combined to cover the whole track over multiple SWRs. Therefore, it appears likely that our simplified model lacks some additional mechanisms that contribute to the termination of individual SWRs in the experiments. For example, building on the work of York and van Rossum (2009), some rate-based models of sequence replay in a circular environment (Romani and Tsodyks, 2015; Theodoni et al., 2018) included short-term synaptic depression as a mechanism for terminating replay. However, Guzman et al. (2016) found pseudo-linear short-term plasticity profiles for the recurrent PC-PC connections at physiological temperatures (although depression was present in recordings at room temperature). Moreover, PCs in our simulations typically fired single spikes at relatively low rates rather than bursts during SWRs (which is similar to the *in vitro* observations of Schlingloff et al. (2014) but in contrast to the *in vivo* results of (Mizuseki and Buzsáki, 2013)), which rules out short-term synaptic plasticity as a key termination mechanism in our model. Another possibility is that an additional cell type that is not currently included in our model is responsible for the termination of SWR events. This explanation was supported by some exploratory simulations where we found that the duration of SWRs could be controlled by a second type of interneuron that provided delayed, long-lasting feedback inhibition to the PCs in the model.

Finally, we have presented our work as a model of the hippocampal CA3 area. This is because this area is known to be able to generate SWRs on its own, has modifiable recurrent connections that are thought to be essential for memory, and there are sufficient experimental data from this region to constrain the model. However, other cortical regions such as CA2, the subiculum, and entorhinal cortex, are likely involved in the initiation of SWRs under normal conditions (Oliva et al., 2016, 2020), and the mechanisms we describe here may be operational in these and other brain areas as well.

### 3.3 Previous models of sharp wave-ripples and sequence replay

In this study, our main objective was to build a simplified, yet powerful model of area CA3 that is strongly constrained by experimental data at all levels, and thus allows us to uncover the mechanistic links between learning, neural population dynamics, and the representation of spatial (or other) sequences in the hippocampus during different behavioral states. Although there is a plethora of hippocampal models that shed light on some of these aspects (these models have been recently reviewed (Buzsáki, 2015; Jahnke et al., 2015) and are also cited when relevant throughout the Results section), there are only a handful of recent models that attempted to treat all of them within a single coherent framework.

The study of Jahnke et al. (2015) is probably the most similar in spirit to ours, as it also explores the relationship between learning, replay, and SWRs. One distinguishing feature of their model is that it relies on the nonlinear amplification of synaptic inputs by dendritic spikes in CA3 PCs for the generation of both sequential activity and ripple oscillations (Memmesheimer, 2010). Importantly, replay always occurs in the forward direction in their model, as it relies on feed-forward chains of strong weights in the network, established by an asymmetric STDP rule that is quite distinct from the one that was later found empirically by Mishra et al. (2016). In addition, the generation of ripple oscillations in their model relies on synchronized pulses of activity generated by interconnected PCs, while recent experimental findings appear to provide causal evidence for the involvement of fast-spiking PVBCs in ripple frequency oscillation generation (Racz et al., 2009; Ellender et al., 2010; English et al., 2014; Schlingloff et al., 2014; Stark et al., 2014; Gulyás and Freund, 2015; Buzsáki, 2015; Gan et al., 2017). Finally, SWRs need to be evoked by synchronous external input in the model of Jahnke et al. (2015), while they can also emerge spontaneously in our model.

Malerba and Bazhenov (2019) developed a combined model of areas CA3 and CA1 to study the generation of sharp waves in CA3 and associated ripple oscillations in CA1. This model relies on distance-dependent connection probabilities in the network for the generation of spatially localized SWR events. The study shows that modifying the recurrent excitatory weights via an asymmetric STDP rule during a simulated learning epoch biases the content of SWRs towards the (forward) reactivation of learned trajectories. Ripple oscillations are modeled only in CA1 and, in contrast to our model (and the models of Taxidis et al. (2012); Donoso et al. (2018); Ramirez-Villegas et al. (2018)), their generation is independent of recurrent inhibition.

A notable recent example of functionally motivated (top-down) modeling of these phenomena is the study of Nicola and Clopath (2019). The authors designed and trained (using supervised learning methods) a network of spiking neurons to generate activity sequences that were either tied to the population-level theta oscillation, or occurred spontaneously in isolation (in a compressed manner), depending on the presence or absence of an external theta-frequency input. Interestingly, these results were achieved by tuning specifically the inhibitory weights in the network, while all other models (including ours) rely on plasticity in the recurrent excitatory synapses. Their model produced forward replay of sequences by default; however, sequences could be reversed by the activation of a distinct, dedicated class of interneurons.

To our best knowledge, ours is the first model that autonomously generates SWRs and replay in a spiking network model using synaptic weights established via the experimentally observed symmetric STDP rule. The model of Haga and Fukai (2018) used symmetric STDP (in combination with short-term plasticity) to modify an existing (pre-wired) weight structure, and showed that these changes biased evoked activity sequences towards reverse replay. Neither spontaneous sharp-waves nor ripple oscillations were observed in this model.

We believe that our approach of fitting the parameters of our single cell models directly to experimental data to mimic the physiological spiking behavior of real PCs and PVBCs is also quite unique. This enabled our models of PCs to capture spike frequency adaptation, which proved to be essential for the generation of propagating activity (sequence replay) despite the essentially symmetric nature of synaptic interactions.

### 3.4 Conclusions

At a more general level, our results highlight the significance of some previously neglected interactions between three fundamental components of brain function: population dynamics, coding, and plasticity. Specifically, the different types of population dynamics (including oscillations and transients such as sharp waves) are mostly seen as a background over which coding and plasticity occur, while the impacts of plasticity and especially neural representations on the generation of population activity patterns are rarely considered. However, our results strongly suggest that the structured network interactions resulting from activity-dependent learning (or development) lead to specific spatio-temporal activity patterns (such as autonomously generated replay sequences), which are in turn critical for the emergence of physiological population activity (such as sharp waves and ripple oscillations). Therefore, our study indicates that the complex structure of synaptic interactions in neuronal networks may have a hitherto unappreciated degree of control over the general mode of activity in the network, and should be taken into account by future theories and models of population activity patterns in any part of the nervous system.

## 4 Methods

In order to investigate the mechanisms underlying hippocampal network dynamics and how these are affected by learning, we built a simplified network model of area CA3. This scaled-down version of CA3 contained 8000 PCs and 150 PVBCs, which is approximately equivalent to cell numbers in area CA3 in a 600 *μm* thick hippocampal slice based on our previous estimates (Schlingloff et al., 2014), which are also in good agreement with other estimates (Bezaire and Soltesz, 2013; Donoso et al., 2018). The connection probability was estimated to be 25% for PVBCs (Schlingloff et al., 2014) and 10% for PCs (Table 3), and was independent of distance in the model. Half of the PCs were assumed to have place fields on the simulated 3 m long linear track.

**Table 2:** Optimized parameters of PC (AdExpIF and ExpIF) and PVBC models. Physical dimensions are as follows: *C_m_*: pF, *g_L_* and *a*: nS, *V_rest_*, Δ*T*, *ϑ*, *θ* and *V_reset_*: mV, *t_ref_* and *τ_w_*: ms, *b*: pA.

| | $C_m$ | $g_L$ | $V_{rest}$ | $\Delta T$ | $\vartheta$ | $\theta$ | $V_{reset}$ | $t_{ref}$ | $\tau_w$ | $a$ | $b$ |
| --- | --- | --- | --- | --- | --- | --- | --- | --- | --- | --- | --- |
| **PC** | 180.13 | 4.31 | -75.19 | 4.23 | -24.42 | -3.25 | -29.74 | 5.96 | 84.93 | -0.27 | 206.84 |
| **PC** | 344.18 | 4.88 | -75.19 | 10.78 | -28.77 | 25.13 | -58.82 | 1.07 | - | - | - |
| **PVBC** | 118.52 | 7.51 | -74.74 | 4.58 | -57.71 | -34.78 | -64.99 | 1.15 | 178.58 | 3.05 | 0.91 |

**Table 3:** Synaptic parameters (taken from the literature or optimized). Physical dimensions are as follows: 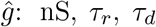 and *t_d_* (synaptic delay): ms and connection probability *p_conn_* is dimensionless. GC stands for the granule cells of the dentate gyrus. (*GC* → *P C* synapses are referred as mossy fibers.)

| | $\hat{g}$ | | $\tau_r$ | $\tau_d$ | $t_d$ | $p_{conn}$ |
| --- | --- | --- | --- | --- | --- | --- |
|  | sym. | asym. |  |  |  |  |
| PC → PC | 0.1-6.3 | 0-15 | 1.3 | 9.5 | 2.2 | 0.1 |
| PC → PVBC | 0.85 | | 1 | 4.1 | 0.9 | 0.1 |
| PVBC → PC | 0.65 | | 0.3 | 3.3 | 1.1 | 0.25 |
| PVBC → PVBC | 5 | | 0.25 | 1.2 | 0.6 | 0.25 |
| GC → PC | 19.15 | 21.5 | 0.65 | 5.4 | - | - |

Only PCs received external input; during exploration, they were activated directly to model place-associated firing; otherwise, they received synaptic input (through the mossy fibers) in the form of uncorrelated Poisson spike trains with a mean rate of 15 Hz. As the hallmark of granule cell activity in the dentate gyrus is sparse, low frequency firing (Jung and McNaughton, 1993; Skaggs et al., 1996), and each CA3 PC is contacted by only a few mossy fibers, the physiological mean rate of mossy fiber input to PCs is probably substantially lower (especially in off-line states of the hippocampus). On the other hand, real CA3 PCs also receive direct input from the entorhinal cortex, and also spontaneous EPSPs from a large population of recurrent collateral synapses. Overall, these other numerous, but small-amplitude inputs may be responsible for most of the depolarization required to bring CA3 PCs close to their firing threshold, in which case mossy fiber input at a significantly lower rate would be sufficient to evoke the same number of action potentials in CA3 PCs.

Network simulations were run in `Brian2` (Stimberg et al., 2019). Learning of the structure of recurrent excitation, single cell and network optimization, and the analysis of the network simulations are detailed in the following sections. A comprehensive list of assumptions made during the model building process is presented in Table 1.

### 4.1 Spike trains during exploration

Spike trains mimicking CA3 PC activity during exploration were generated with exponentially distributed inter spike intervals (ISIs) with mean 1*/λ*, giving rise to Poisson processes. Spike trains of non-place cells had mean firing rates of *λ* = 0.1 Hz. For the spike trains of the randomly selected 4000 place cells, homogeneous Poisson processes with *λ* = 20 Hz were generated, and spike times were accept-reject sampled with acceptance probability coming from place cell-like tuning curves (eq. (2)), which led to inhomogeneous Poisson processes with time-dependent rate *λ*(*t*). Tuning curves were modeled as Gaussians centered at randomly distributed positions and standard deviation set to cover 10% of the 3 m long linear track (eq. (1)). Edges of the place fields were defined where the firing rate dropped to 10% of the maximal 20 Hz (Dragoi and Buzsáki, 2006). Tuning curves were also modulated by the background *f_θ_* = 7 Hz theta activity, phase precessed up to 180° at the beginning of the place field (O’Keefe and Recce, 1993). The firing rate of the *i*th place cell was calculated as follows:

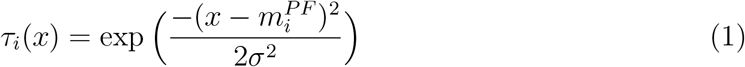

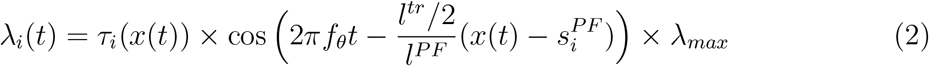

where *τ_i_*(*x*) is the spatial tuning curve of the *i*th neuron, *x*(*t*) is the position of the animal, 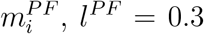 and 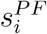 are the middle, length and start of the given place field respectively, while *l ^tr^* = 3 m is the length of the linear track; *λ _max_* = 20*Hz* is the maximum in-field firing rate.

Spikes within a 5 ms refractory period of the previous spike were always rejected. The speed of the animal was set to 32.5 cm/s, thus each run took ~9.2 s, after which the animal was immediately “teleported back” to the start of the linear track. Generated spike trains were 400 s long, leading to ∼ 43 repetitions on the same linear track.

### 4.2 Learning via STDP

STDP was implemented by an additive pair-based learning rule, evaluated at spike arrivals (Kempter et al., 1999; Gerstner et al., 2014). Synaptic weights evolved as follows:

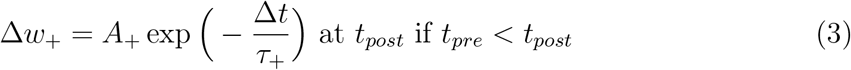

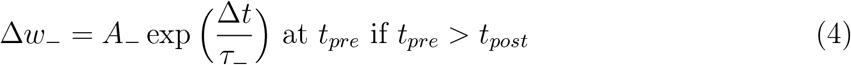

where Δ*t* = *t_post_* − *t_pre_* is the time difference between action potentials, *A_±_* describe the weight update, which decayed exponentially with time constants *τ_±_*. Synaptic weights were cropped at *w_max_* = 20 nS. To reproduce the broad STDP curve presented in Mishra et al. (2016) *τ_±_* = 62.5 ms was used. In the classical asymmetric STDP rules *A*_+_ is positive, while *A_−_* is negative; here, both of them were set to 80 pA to obtain a symmetric STDP curve (Mishra et al., 2016). In simulations using the asymmetric STDP rule, *τ_±_* = 20 ms, *A*_+_ = 400 pA, *A_−_* = −400 pA, and *w_max_* = 40 nS were used. In both cases PCs were sparsely connected (Table 3) and weights were initialized to 0.1 nS. In the learning phase the intrinsic dynamics of the PCs were not modeled explicitly, since only the timing of their spikes mattered, which was set directly as described above. No self-connections were allowed, and diagonal elements of the learned recurrent weight matrix were always set to zero after any modification.

### 4.3 In vitro electrophysiology

Somatic whole-cell patch-clamp recordings were performed in acute hippocampal slices as described before (Papp et al., 2013; Schlingloff et al., 2014; Kohus et al., 2016). Pyramidal cells were recorded in the CA3 pyramidal cell layer of juvenile control mice, while PVBCs were recorded in a targeted manner in transgenic mice that expressed enhanced green flurenscent protein controlled by the parvalbumin promoter (BAC-PV-eGFP) (Meyer et al., 2002). To characterize the physiological response properties of the neurons, hyperpolarizing and depolarizing current steps of various amplitudes were injected into the soma, and the voltage response of the cell was recorded. Injected current pulses had a duration of 800 ms, and amplitudes between −100 and 600 pA. Experimental traces were corrected for the theoretical liquid junction potential before further use.

### 4.4 Single cell models

Neurons were modeled with the AdExpIF model (Naud et al., 2008; Gerstner et al., 2014). AdExpIF neurons are described by their membrane potential *V* (*t*) and the adaptation variable *w*(*t*), which obey:

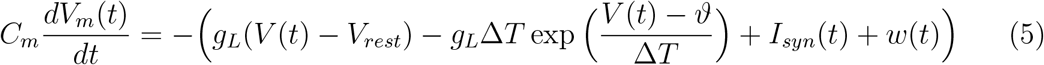

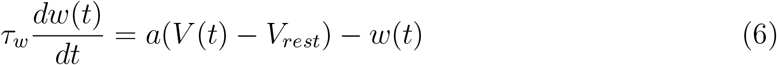

where *C_m_* is the membrane capacitance, *g_L_* is the leak conductance, *V_rest_* is the reversal potential of the linear leak current (which is approximately equal to the resting potential), *ϑ* is the intrinsic spike threshold, Δ*T* characterizes the “sharpness” of the threshold, *w*(*t*) is the adaptation current and *I_syn_* is the synaptic current (see below). When *V* (*t*) crosses the firing threshold *θ*, it is reset to *V_reset_* and remains there for a refractory period *t_ref_*. The adaptation current is also increased by a factor *b* at each spike arrival and decays exponentially afterwards with the time constant *τ_w_*. The parameter *a* describes the strength of sub-threshold adaptation.

To investigate the role of adaptation, an ExpIF PC model was also fit to the data. The ExpIF model is the same as eq. (5) without the *w*(*t*) adaptation current (implemented as an AdExpIF model with parameters *a* and *b* set identically to zero). The parameters of all models were fit to experimental data from our somatic whole-cell recordings, and the voltage responses to current injections of four different amplitudes (including two subthreshold and two suprathreshold stimuli) were used in each case. Parameters were tuned using the `Optimizer` package (Friedrich et al., 2014) with the `NEST` simulator as backend (Gewaltig and Diesmann, 2007). Spike count, ISI distribution, latency to first spike and mean squared error (excluding spikes) were used as equally weighted features. After comparing different optimization techniques, the final parameters presented here were obtained with an evolutionary algorithm implemented by the `inspyred` package (Garrett, 2012), running for 100 generations with a population size of 100. The parameters which yield the best models for the CA3 populations are summarized in Table 2.

### 4.5 Synapse models

Synapses were modeled as conductances with bi-exponential kinetics:

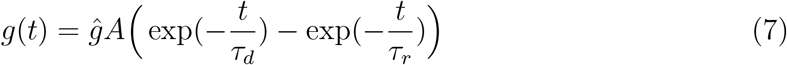

where 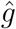 is the peak conductance (which will also be referred to as the synaptic weight) and *τ_r_* and *τ_d_* are rise and decay time constants respectively. The normalization constant 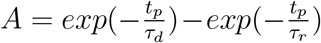 was chosen such that the synapses reach their peak conductance at *t_p_* = *τ_d_τ_r_/*(*τ_d_ − τ_r_*)*log*(*τ_d_/τ_r_*) ms. Kinetic parameters were taken from the literature (Geiger et al., 1997; Bartos et al., 2002; Lee et al., 2014; Vyleta et al., 2016; Guzman et al., 2016) and are summarized in Table 3. The postsynaptic current contained AMPA receptor- and GABA-A receptor-mediated components, and was computed as:

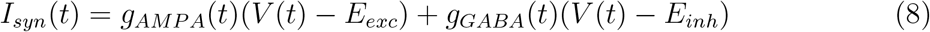

where *E_exc_* = 0 mV and *E_inh_* = − 70 mV are the reversal potentials of excitatory and inhibitory currents, respectively.

### 4.6 Network optimization

Synaptic weights of the network (5 parameters in total) were optimized with an evolutionary algorithm using a custom written evaluator in `BluePyOpt` (Van Geit et al., 2016). The multi-objective fitness function 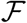, designed for this network included 6 separately weighted features (eq. (9)): physiological PC firing rate, no significant gamma oscillation in the PC population, significant ripple frequency oscillations in the rates of PC and PVBC populations as well as high ripple vs. gamma power in the rates of the PC and PVBC populations:

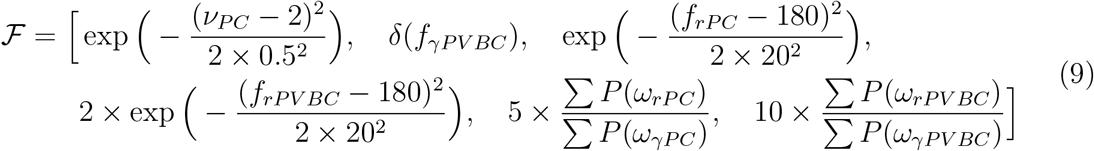

where *ν* is the firing rate, *δ* is the Dirac-delta function, *f_r_* and *f_γ_* are significant peaks in the ripple and gamma range (see below) of the PSD of the firing rate respectively and *P* (*ω_r_*) and *P* (*ω_γ_*) are the periodogram values within the gamma and ripple bands of the firing rate respectively, while the sum of them represents the power within the frequency bands (as below). As in the case of the network simulations (see above) if the PC firing rate exceeded the 2 Hz high activity state detection threshold spectral features were only extracted in these time windows. Sequence replay was not analysed during the optimization. Optimizations were run with 50 offspring for 10 generations. Synaptic weights were varied in the [0.1, 5] nS range, except the “detonator” mossy fiber ones which were given a higher [15, 30] nS range (Henze et al., 2002; Vyleta et al., 2016). For the learned recurrent excitatory weights an additional scaling factor was introduced. All learned weights are presented with this optimized scale factor (0.62 for symmetric and 1.27 for asymmetric STDP rule) taken into account. Final weights are presented in Table 3. The ExpIF PC model required much higher synaptic drive to fire at the same frequency as the AdExpIF model, thus the mossy fiber i nput w eight w as d oubled (38.3 n S) when ExpIF PC models were used.

### 4.7 LFP estimate

An estimate of the LFP was calculated by summing the synaptic currents of a small randomly selected subset of *N* = 400 PCs (Mazzoni et al., 2008). This approach is essentially equivalent to using “transmembrane” currents to calculate the field potential at an arbitrary sampling point *x_e_*, using volume conduction theory and the forward model (Einevoll et al., 2013):

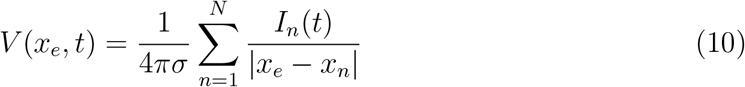

where *σ* = 1/3.54 S/m is the extracellular conductivity and *I_n_*(*t*) denotes the trans-membrane currents of the *n*th neuron. There was no attempt to replicate the spatial organization of CA3 PCs and a uniform |*x_e_ − x_n_*| = 1*μm* distance from the sampling point was used (note that this choice affects the results only as a constant scaling factor). The resulting signal was low pass filtered at 500 Hz with a 3rd order Butterworth filter.

### 4.8 Spectral analysis

Power spectral density (PSD) was estimated by Welch’s method with a Hanning window, using 512 long segments in case of population rates (sampling frequency = 1 kHz) and 4096 long segments for LFP (see below, sampling frequency = 10 kHz) with 0.5 overlap. If the network showed multiple sequence replays during the 10-seconds long simulations (most cases) only the detected high activity states (see above) were analysed and averaged, to get rid of the high power at ~1.5 Hz, signaling the frequency of sequence replays. In this case shorter segments (256 and 2048 respectively) were used to estimate the PSD. The significance of peaks in the power spectra in the gamma (30-100 Hz) and ripple (150-220 Hz) bands was evaluated using Fisher’s g-statistic (Fisher, 1929) defined as:

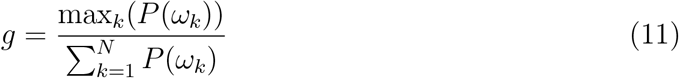

where *P* (*ω*) is the periodogram (Welch’s method estimates PSD by averaging periodograms from the short segments) evaluated at *k* discrete frequencies, *N* is the length of the periodogram and 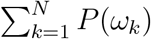 is the total power of the spectrum. The distribution of g-statistics under the null hypothesis (*H*_0_) (Gaussian white noise) is given by:

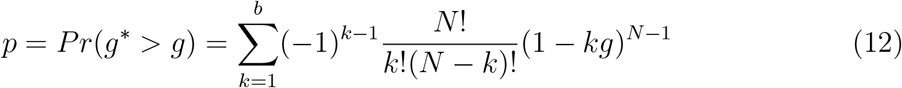

where *b* is the largest integer less than 1*/g*. Large value of *g* (small p-value) indicates a strong periodic component and leads to the rejection of *H*_0_. Alpha level 0.05 was used throughout the study. To characterize non-significant oscillations too, gamma and ripple power (defined as the sum in the given frequency band divided by the total power in the 0-500 Hz range) were calculated as well. Time-frequency representations were created by convolving the firing rates (or LFP) with samples of the integral of the Morlet wavelet Ψ(*t*) = *exp*(*t*^2^/2)*cos*(5*t*) evaluated at the scales corresponding to the 25-325 Hz band, using the pywt package (Lee et al., 2006).

### 4.9 Replay analysis

Sequence replay was analysed with methods used by experimentalists having access to spike times of hundreds of identified neurons (Ólafsdóttir et al., 2018). Firstly, candidate replay events were selected based on the averaged (into 20 ms bins) PC population firing rate crossing the threshold of 2 Hz for at least 260 ms. Secondly, the animal’s position was estimated with a memoryless Bayesian place decoder based on the observed spikes within the selected time windows (Davidson et al., 2009; Karlsson and Frank, 2009). Only spikes from the *N* = 4000 place cells were used. For numerical stability log likelihoods were calculated:

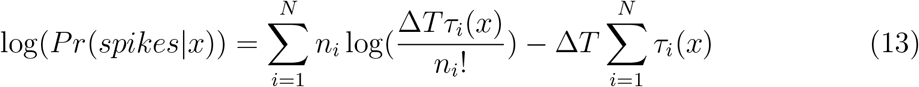

where *n_i_* is the number of spikes of the *i*th neuron within the Δ*T* = 10 ms long, non-overlapping time bins and *τ_i_*(*x*) is the tuning curve used for spike train generation (eq. (1)). The 3 m long linear track was binned into 50 intervals, resulting in 6 cm spatial resolution. Thirdly, constant velocity *v* neural trajectories were detected with a 2D band finding method in the decoded posterior matrix (Davidson et al., 2009). For candidate events consisting of *n* time bins, the average likelihood *R* that the animal is within distance *d* = 18 cm of a particular trajectory is given by:

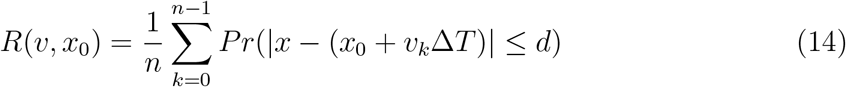

where *x*_0_ is the starting position of the trajectory. *R*(*v, x*_0_) was maximized using an exhaustive search to test all combinations of *v* between −18 m/s and 18 m/s in 0.3 ms/s increments (excluding slow trajectories with speed ∈ [−0.3, 0.3] m/s) and *x*_0_ between −1.5 m and 4.5 m in 3 cm increments. Lastly, to determine the significance of replay, *R_max_* was compared to the best line fits of 100 posterior probability matrices generated by shuffling the identities of cells included in the candidate event. Only events with *R_max_* values exceeding the 95th percentile of their own shuffled distribution were labeled as replay.

To replicate the step-size analysis of Pfeiffer and Foster (2015) the position of the animal was estimated as a weighted average based on the posterior matrix in each time bin instead of the band finding method. As their control distribution for the skewed step-sizes (“predicted step-size” distribution) was derived for a 2D arena, it was not directly applicable to our linear track setup. Therefore, we defined the predicted step-size distribution based on the ratio of the length of the replayed path and the duration of the replay event for the SWRs detected in the simulations.

## Accessibility

The source code is publicly available at https://github.com/KaliLab/ca3net.

## Acknowledgements

We thank G. Buzsáki, D. Csordás, G. Nyíri, B. Ujfalussy, and V. Varga for insightful comments on the manuscript, M. Stimberg for his help with Brian2, and F. Crameri for “scientific” colormaps.

## Author contribution

S.K., A.G. and T.F. conceptualized the study. S.K. supervised the study. M.R.K., O.I.P., N.H., and A.G. provided experimental data. A.E., E.V., O.N. and S.K. worked on spike train generation and learning. B.B. and S.K. optimized single cell models. A.E., B.B., E.V. and S.K. built and optimized the network and analysed the results. A.E., I.M. and S.K. designed and implemented weight matrix modifications. A.E. created all figures. A.E. and S.K. wrote the manuscript with inputs from all authors.

---

**Figure S1:**
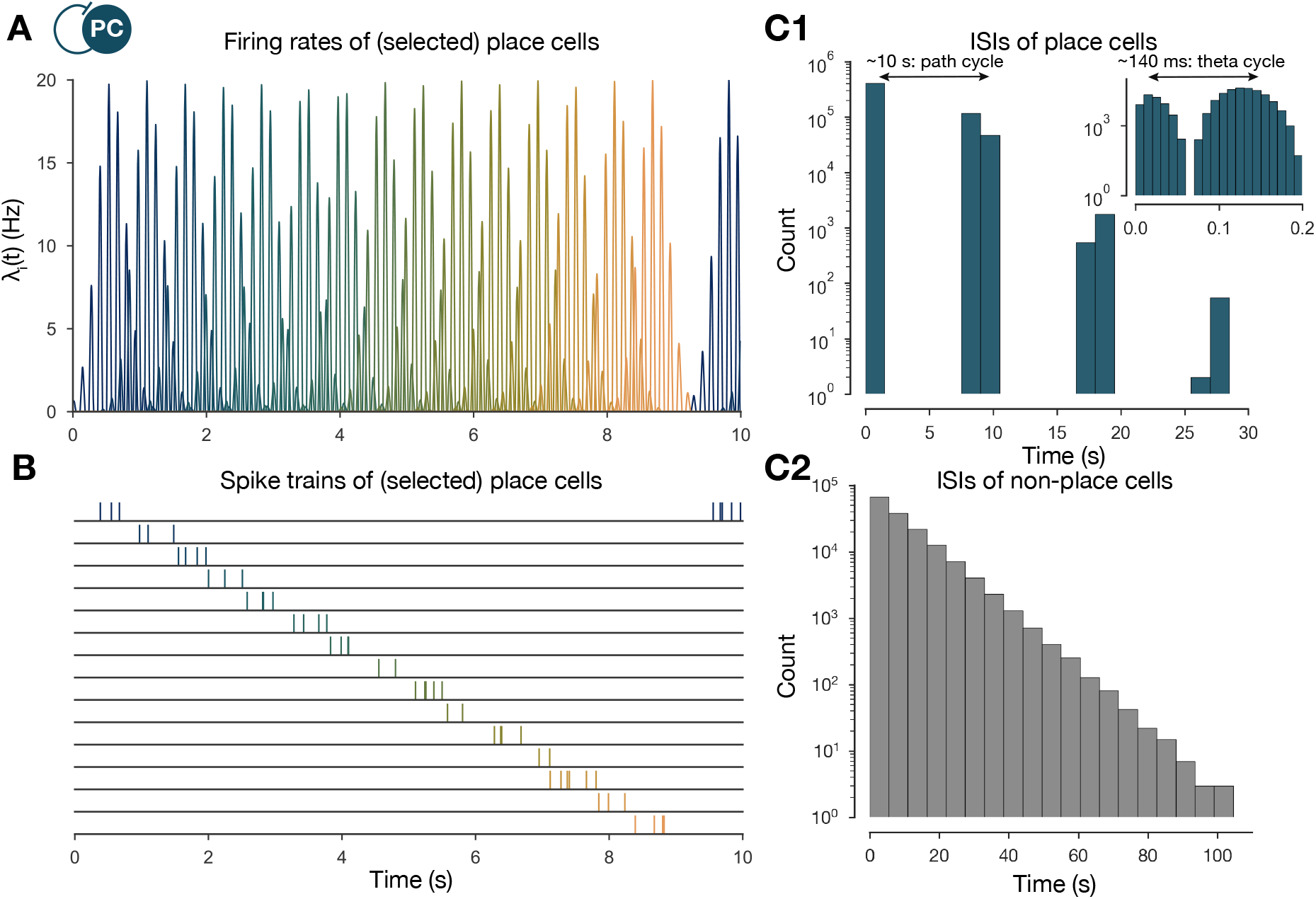
The generation of the spike trains of PCs in the exploration phase. **(A)** Firing rates of exemplar place cells covering the whole 3 m long linear track. Compared to the tuning curves shown in Figure 1A (eq. (1)), these are time-dependent rates modulated by theta oscillation and phase precession (eq. (2)). **(B)** Exemplar spike trains generated based on the firing rates shown in (A). (Spike trains used in the learning phase were 400 second long. For the purpose of visualization, only the beginning is shown here.) **(C)** ISI distribution of the generated spike trains. ISIs of place cells (C1) (insert is a zoom into the same distribution at a finer timescale to show theta modulation) and non-place cells (C2).

**Figure S2:**
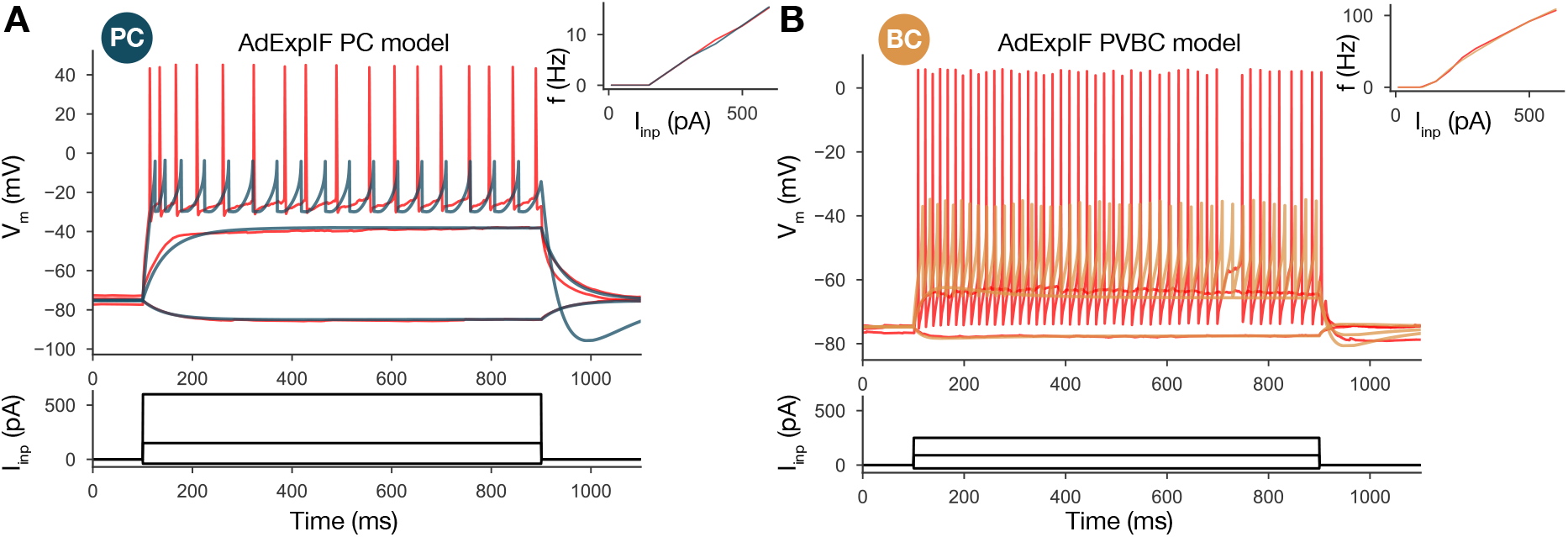
Single cell models. **(A)** Fitted AdExpIF PC model (blue) and experimental traces (red) are shown on the top panel. The 800 ms long step current injections shown in the bottom were as follows: −0.04, 0.15 and 0.6 nA. **(B)** Fitted ExpIF PVBC model (gold) and experimental traces (red) are shown in the top panel. The amplitudes of the 800 ms long step current injections shown at the bottom were as follows: −0.03, 0.09 and 0.25 nA. Inserts show the fI curve of the *in vitro* (red) and *in silico* cells. For parameters of the cell models see Table 2.

**Figure S3:**
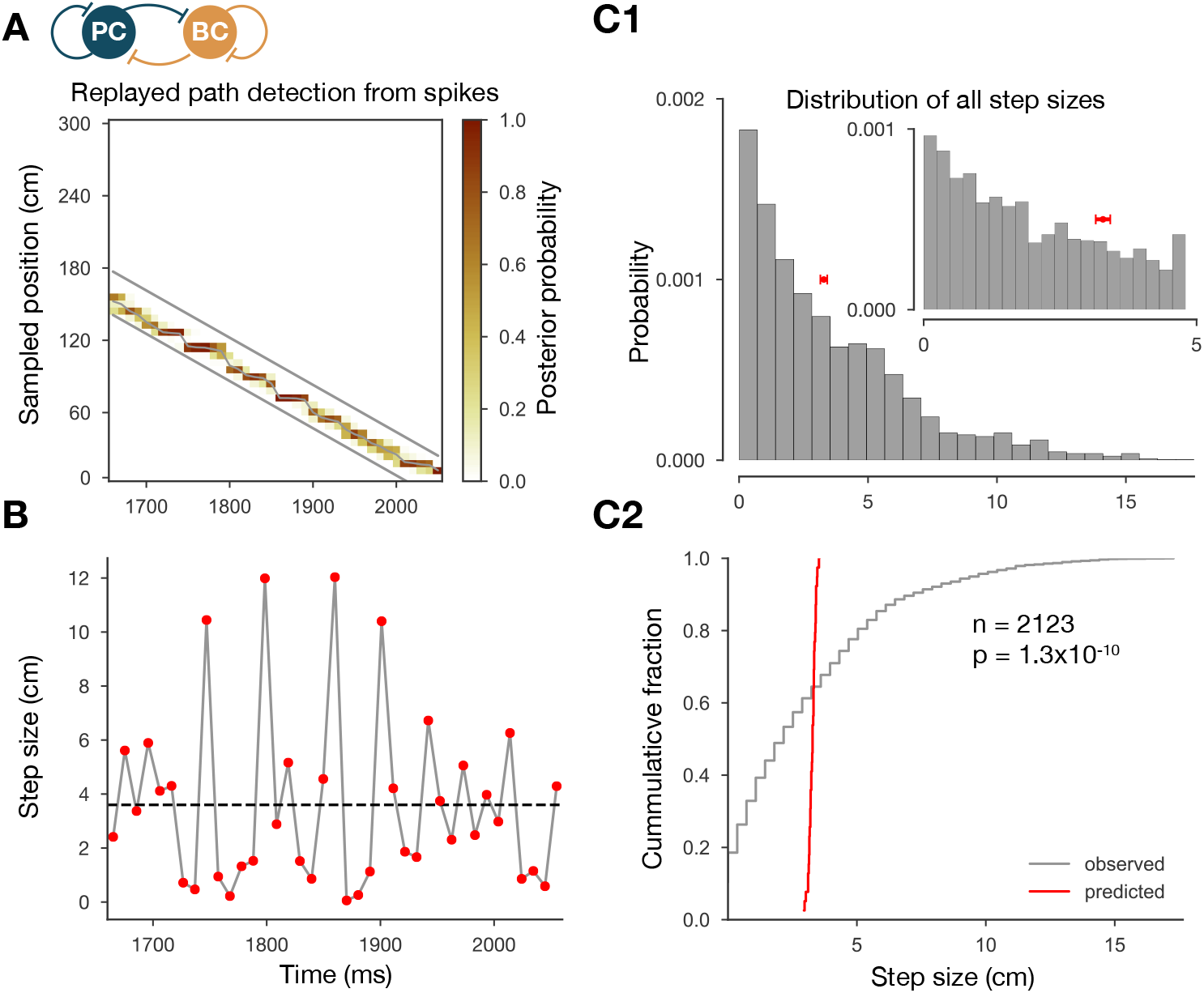
The step-size distribution of the decoded paths is much wider than expected. **(A)** Posterior matrix of the decoded positions from spikes within a selected high activity state (1st one from Figure 2A). Thick gray lines indicate the edges of the decoded, constant velocity path. Thin gray line shows the decoded path by connecting the weighted average positions in every 10 ms long time step. **(B)** Step sizes from the decoded, variable velocity path (see (A)) for the same period (1st high activity state in Figure 2A). The horizontal dashed black line shows the average or predicted step-size within the given period. **(C1)** Skewed distribution of observed step sizes (in gray) and the predicted (from evenly spacing) step-size distribution (in red) for more than a hundred replay events (similar to the one in (A) and (B)). Inserat shows the same distributions at smaller scales. **(C2)** Cumulative distribution of the observed and predicted step sizes shown in (C1). Observed vs. predicted distributions significantly differ (two-sample Kolmogorov-Smirnov test).

**Figure S4:**
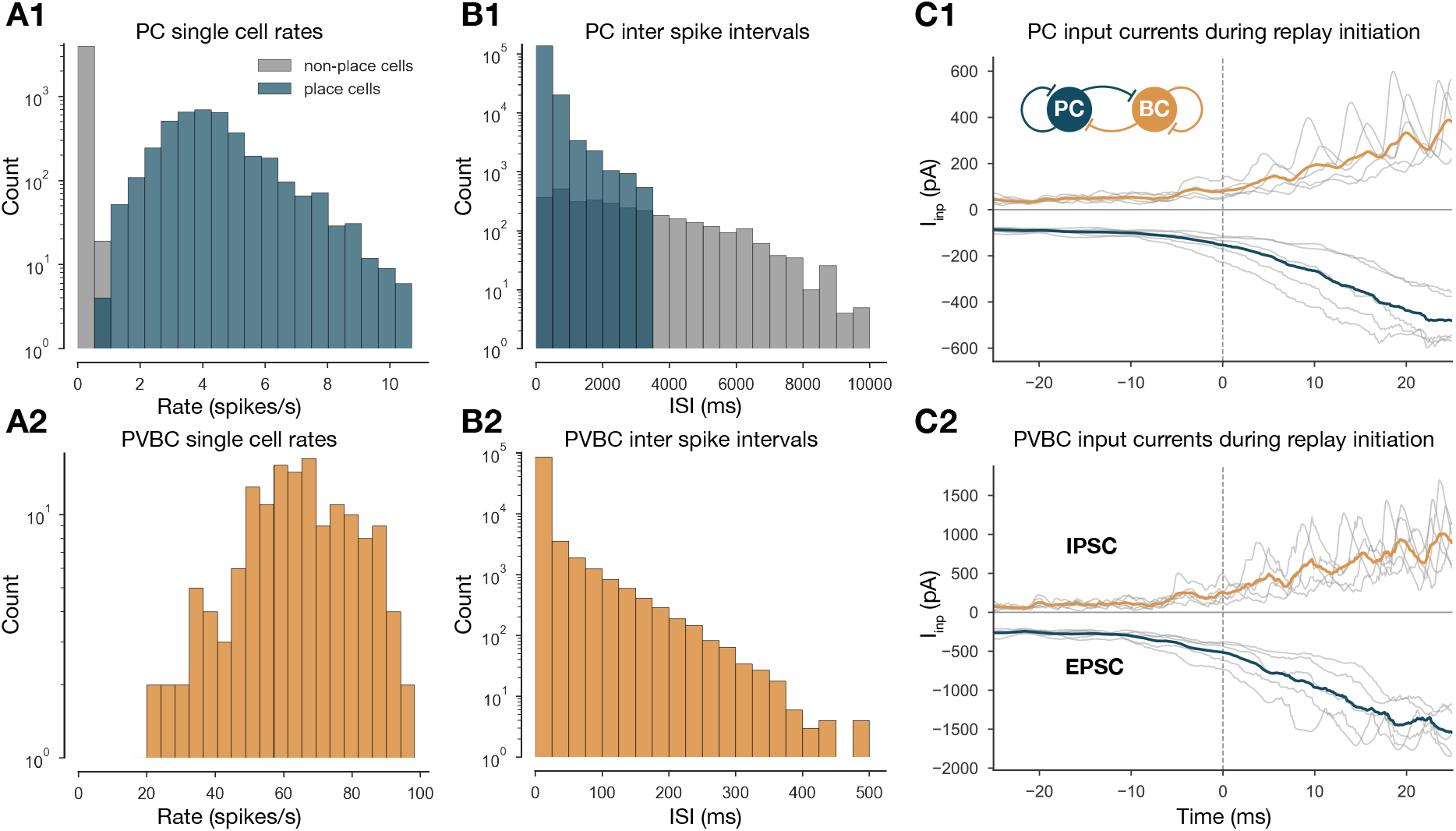
Single cell characteristics during network simulations. **(A)** Single cell PC (A1) and PVBC (A2) firing rates of the 10 second long simulation shown in Figure 2. **(B)** ISIs of PCs (B1) and PVBCs (B2) in the 10-second long simulation shown in Figure 2. **(C)** Synaptic input currents of PCs (C1) and PVBCs (C2) during sequence replay initiation. Gray lines are the averages of the EPSCs and IPSCs of 400 PCs and 30 PVBCs respectively. Individual gray lines correspond to individual high activity states (n=7, see Figure 2A). Dashed vertical lines (at 0 ms) indicate the beginning of the periods marked as high activity states (see Figure 2A). Colored lines represent the grand average EPSC (blue) and IPSC (gold) arriving at PCs (C1) and PVBCs (C2) during SWR initiation.

**Figure S5:**
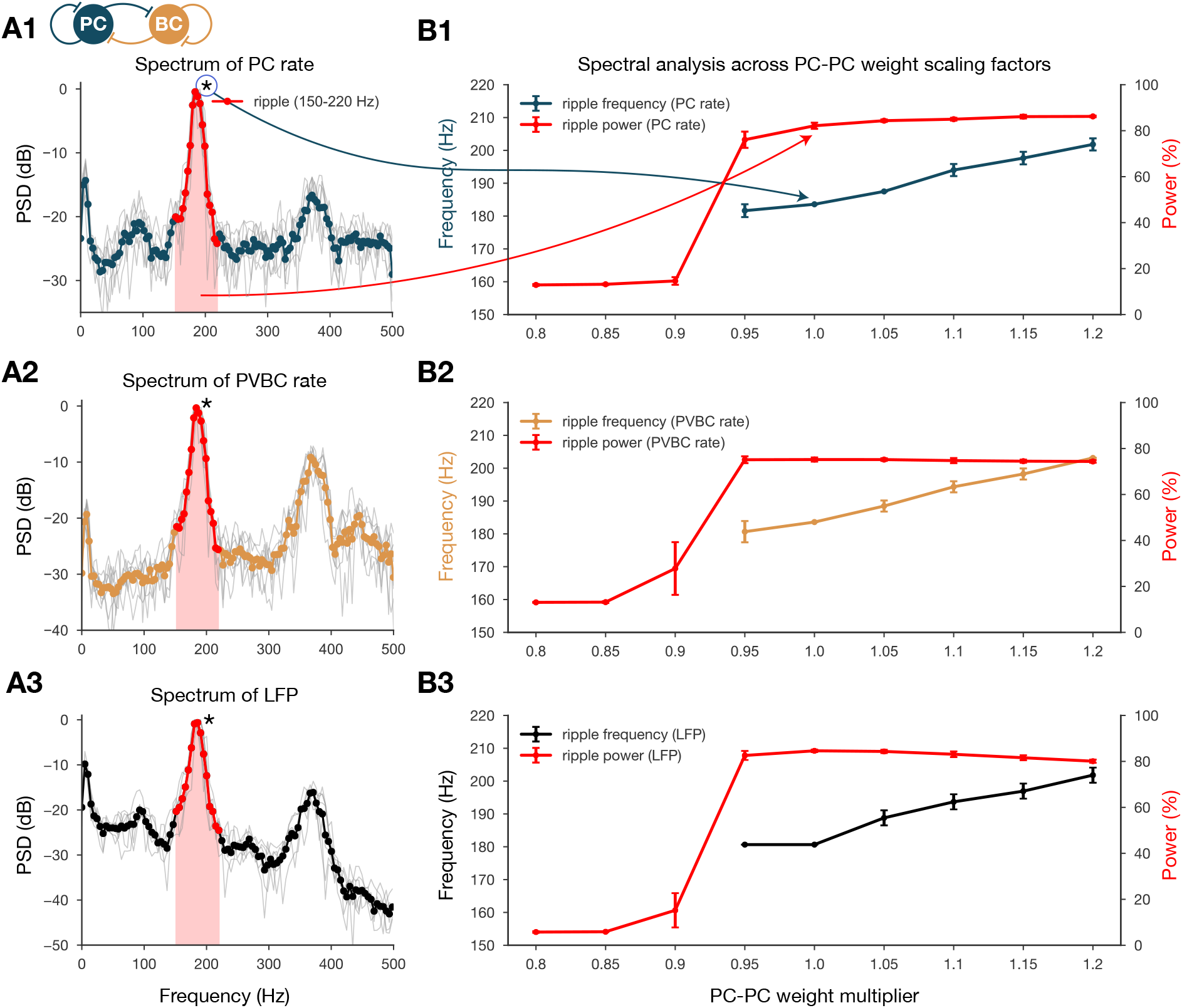
Spectral analysis of network dynamics across different PC-PC weight scaling factors. **(A)** PSDs of PC (A1) and PVBC (A2) population rates and estimated LFP (A3). Gray lines correspond to individual high activity states (n=14) shown in Figure 2A, while the thicker colored lines are their averages. Ripple frequency range (150-220 Hz) is highlighted in red. Shaded red area below the curves indicates the power in the ripple range. **(B)** Spectral analysis of network dynamics across different E-E weight scaling factors (0.8-1.2). The frequency of significant ripple oscillation and ripple oscillation power (red) are shown for PC (B1) and PVBC (B2) population rates and estimated LFP (B3). (B3) is the same as the bottom panel of Figure 3B and it is duplicated only to show how similar the curves are for the rates and the estimated LFP.

